# A lncRNA/Lin28/*Mirlet7* axis coupled to DNA methylation fine tunes the dynamics of a cell state transition

**DOI:** 10.1101/131110

**Authors:** Meng Amy Li, Paulo P. Amaral, Priscilla Cheung, Jan H. Bergmann, Masaki Kinoshita, Tüzer Kalkan, Meryem Ralser, Sam Robson, Ferdinand von Meyenn, Maike Paramor, Fengtang Yang, Caifu Chen, Jennifer Nichols, David L. Spector, Tony Kouzarides, Lin He, Austin Smith

## Abstract

Execution of pluripotency requires progression from the naïve status represented by mouse embryonic stem cells (ESCs) to a condition poised for lineage specification. This process is controlled at transcriptional, post-transcriptional and epigenetic levels and non-coding RNAs are contributors to this regulation complexity. Here we identify a molecular cascade initiated by a long non-coding RNA (lncRNA), *Ephemeron* (*Epn*), that modulates the dynamics of exit from naïve pluripotency. *Epn* deletion delays the extinction of ESC identity, an effect mediated by perduring expression of the pivotal transcription factor Nanog. In the absence of *Epn*, Lin28a expression is reduced, resulting in an elevated level of *Mirlet7g* that suppresses *de novo* methyltransferases Dnmt3a/b. *Dnmt3a/b* deletion also retards exit from the ESC state, and is associated with delayed promoter methylation and slower down-regulation of *Nanog.* Altogether, our findings reveal a lncRNA/miRNA/DNA methylation axis that facilitates a timely stem cell state transition.

## Introduction

Mouse embryonic stem cells (ESCs), *in vitro* counterparts of the pre-implantation epiblast, exhibit dual properties of self-renewal and differentiation (Boroviak et al., 2015; Bradley et al., 1984; Evans and Kaufman, 1981; Martin, 1981). These properties make them an attractive system for investigating cell fate decision making. In the embryo, spatially and temporally coordinated signals direct the rapid and continuous transition of the epiblast towards lineage specification (Acampora et al., 2016; Kalkan et al., 2017). In contrast, ESCs can be suspended in a ground state of pluripotency, where self-renewal is decoupled from lineage specification, using two inhibitors (2i) of glycogen synthase kinase 3 (GSK3) and mitogen-activated protein kinase kinase (MEK1/2), along with cytokine leukaemia inhibitory factor (LIF) (Ying et al., 2008). Therefore, ESCs provides a unique opportunity to explore the principles and molecular players underlying the developmental progression of pluripotency (Kalkan and Smith, 2014).

While it is increasingly clear that the ESC state is maintained by a core network of transcription factors (Dunn et al., 2014; Ivanova et al., 2006; Niwa et al., 2000; Niwa et al., 2009; Wray et al., 2011), little is known about how cells progress from this state to lineage specification (Buecker et al., 2014; Kalkan and Smith, 2014). Loss-of-function screens have highlighted a multi-layered machinery for dissolving this transcription factor network (Betschinger et al., 2013; Leeb et al., 2014) and the latency period for exiting the naïve state depends on the clearance kinetics of the network members (Dunn et al., 2014). The coordination of multiple antagonistic regulators thus ensures a rapid and complete dismantling of this core network and consequent timely extinction of ESC identity upon 2i withdrawal (Kalkan and Smith, 2014).

In addition to protein coding genes, accumulating evidence suggests that non-coding RNAs can contribute to the regulation complexity for cell fate transitions. Within this class, long non-coding RNAs (lncRNAs) comprise a large fraction of the transcriptome in diverse cell types and exhibit specific spatio-temporal expression (FANTOMConsortium, 2005; Guttman et al., 2009; Necsulea et al., 2014). The genomic distribution of lncRNAs is non-random (Luo et al., 2016) and a subclass of lncRNAs which are divergently transcribed from the neighbouring genes, are thought to regulate proximal gene expression *in cis*, either due to the process of transcription (Ebisuya et al., 2008; Engreitz et al., 2016; Martens et al., 2004) or through lncRNA-protein interactions to recruit regulatory complexes (Lai et al., 2013; Lee, 2012; Luo et al., 2016; Nagano et al., 2008). However, the functions and mode of action of the vast majority of lncRNAs remain unknown and require case-by-case experimental investigation. In mouse ESCs, knockdowns of a number of lncRNAs have been reported to exert effects on the transcriptome (Bergmann et al., 2015; Dinger et al., 2008; Guttman et al., 2011; Lin et al., 2014; Sheik Mohamed et al., 2010) and in some cases impair self-renewal (Lin et al., 2014; Luo et al., 2016; Savić et al., 2014). Therefore, lncRNAs could provide an additional layer of regulation in cell fate transition from the ESC state.

We investigated the potential involvement of lncRNAs in transition from the naïve ESC state and identified a dynamically regulated lncRNA that we named *Ephemeron* (*Epn*). We present functional evaluation of *Epn* and delineation of a downstream molecular cascade, which is an integral part of the regulatory machinery driving the irreversible exit from naïve pluripotency.

## Results

### Identification of lncRNAs associated with transition from naïve pluripotency

Post-implantation epiblast derived stem cells (EpiSCs) represent a primed state of pluripotency developmentally downstream of the naïve state ESCs (Brons et al., 2007; Nichols and Smith, 2009; Tesar et al., 2007). To identify lncRNA candidates with a possible role in ESC transition, we analysed *in silico* the effect of genetic perturbation on expression of ESC and EpiSC states based on published data. We first selected genes that are over ten-fold differentially enriched in ESCs (182 genes) and EpiSCs (131 genes) relative to each other (Tesar et al., 2007) as molecular signatures to represent these two states. We investigated the impact on these two signature sets when individual lncRNAs (147 in total) and known protein coding regulators (40 in total) were knocked down in ESCs grown in LIF/serum based a published study (Guttman et al., 2011) (Fig1A, Fig1-source data 1). Serum culture supports a heterogeneous mixture of naïve and primed cells (Chambers et al., 2007; Kolodziejczyk et al., 2015; Marks et al., 2012). Therefore, analysis in this condition could potentially reveal regulators of the ESC and EpiSC states. The effect of each gene knockdown was plotted based on the percentage of genes significantly altered within ESC and EpiSC signature sets (FDR<0.05 and fold change > 2 or <0.5 over negative control defined by the original study). Applying this approach, we identified lncRNAs that increased ESC and decreased EpiSC signatures when knocked down, suggestive of a potential role in transition from the ESC state (Fig1A bottom right quadrant). We validated the approach by analysing the knockdown effects of known ESC self-renewal regulators. As predicted, depletion of factors that maintain the ESC state, such as Stat3, Esrrb, Sox2 and Klf4, led to a decrease in ESC and increase in EpiSC signature (Fig1A), while knockdown of Oct4 gave rise to a decrease in both ESC and EpiSC signatures consistent with its requirement in both states (Niwa et al., 2000; Osorno et al., 2012).

**Figure 1:**
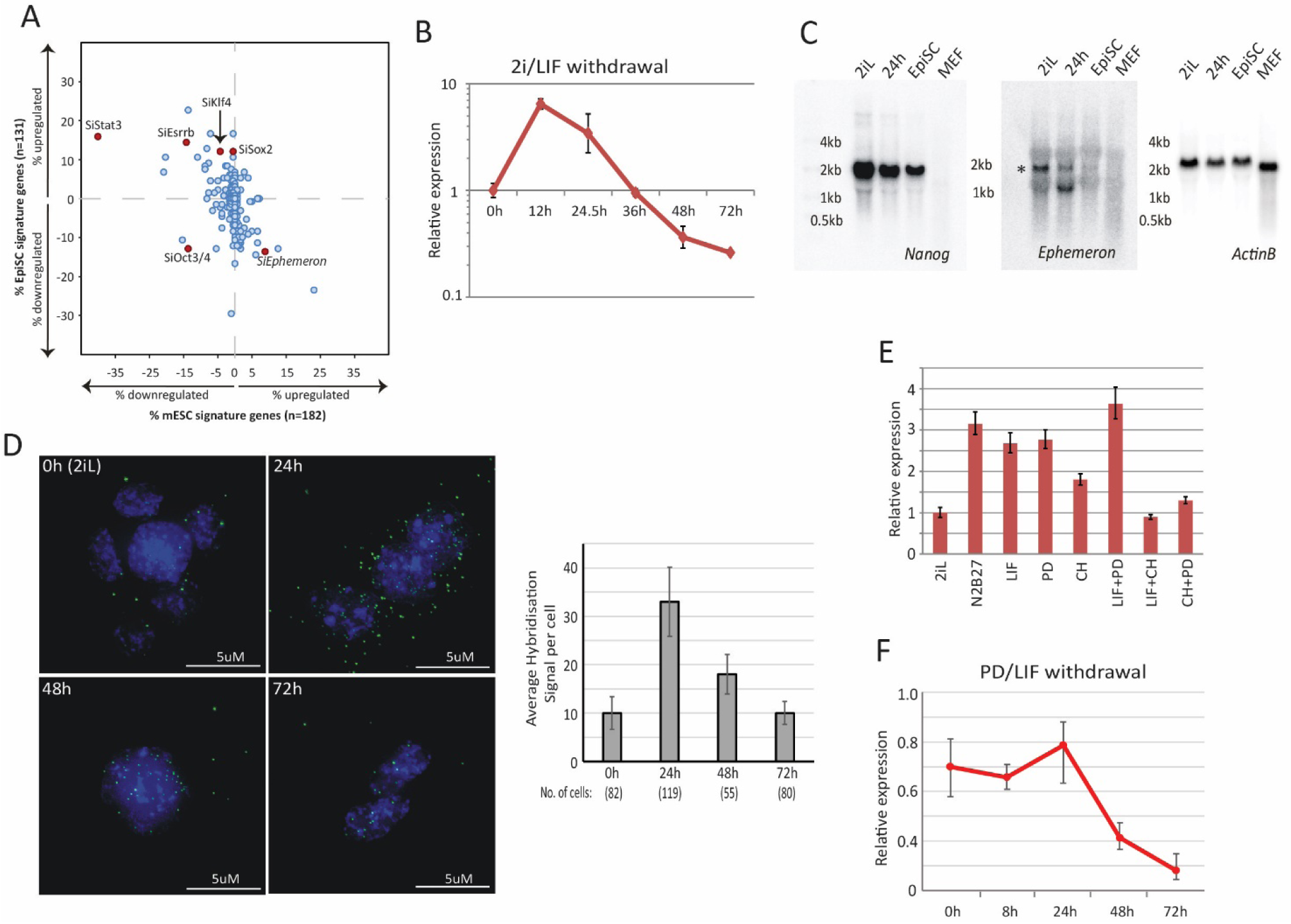
Dynamic expression of lncRNA *Ephemeron* during exit from naïve pluripotency. A, Bioinformatic analysis of potential lncRNA candidates in naïve state regulation based on published transcriptome data for lncRNA and pluripotency related gene knockdowns. Each dot represents the effect on ESC (x-axis) and EpiSC (y-axis) gene signatures when a given gene is knocked down. B, RT-qPCR detection of *Epn* expression relative to *β-actin* upon 2iL withdrawal. Mean+/-SD, n=3. C, Northern blotting of *Epn*, *Nanog* and *β-actin* in ESCs in 2i/LIF or withdrawn from 2i/LIF for 24 hours, EpiSCs and MEF. * indicates a cross hybridising RNA species since part of the probe region overlaps with TEs. D, RNA-FISH for *Epn* upon 2i/LIF withdrawal with quantification of average hybridisation signals per cell. Mean value of total hybridisation signals for all cells +/-SD, n=2. E, *Epn* expression relative to *β-actin* upon 2i/LIF component withdrawal quantified by RT-qPCR. Cells cultured in 2i/LIF and were transferred to N2B27 containing indicated single or dual factors for 24 hours. Mean+/-SD, n=3. F, *Epn* expression relative to *β-actin* upon PD/LIF withdrawal quantified by RT-qPCR. Mean+/-SD, n=3.

We next examined expression profiles of these candidate lncRNAs during exit from self-renewal in defined conditions, exploiting the Rex1::GFP (RGd2) reporter ESC cell line (Kalkan et al., 2017; Wray et al., 2011) (Fig1– source data 2). Upon 24 hours post 2i withdrawal, Rex1 expression status can discriminate subpopulations of cells with distinct functional properties, with Rex1-GFP high cells corresponding to undifferentiated ESCs and low cells corresponding to cells with extinguished ESC identity (Kalkan et al., 2017). Amongst the 16 candidates analysed, *linc1281* (Refseq entry D630045M09Rik) (Fig1 – Figure Supplement 1A) was the third highest expressed lncRNA across all time points. Notably this lncRNA showed a distinctive profile during the first 24 hours, with differential expression observed between Rex1-GFP high and low cells (Fig1B, Fig1- Figure Supplement 1B). Due to its dynamic and transient expression profile, we chose the name *Ephemeron* (*Epn*). Ribosomal profiling analysis indicated that *Epn* is indeed a non-coding RNA, with the longest predicted open reading frame (80 amino acids) possessing a ribosome release score typical of a non-coding sequence (Guttman et al., 2013). *Epn* is located in a region of high transposable element (TE) content, with its exons comprised of 76.4% annotated TE sequences (including ERV-K, LINE L1, and SINE B2 elements, Fig1- Figure Supplement 1A). This genomic region exhibits minimal sequence conservation in mammals (Fig1 – Figure Supplement 1A) and we failed to identify any human homologue either within the syntenic region or elsewhere in the human genome. However, a positionally conserved spliced transcript (CA504619) that shares 79% sequence identity to exon 3 of mouse *Epn* is present within the rat syntenic region (Fig1 – Figure Supplement 1C). Therefore, it is likely that *Epn* is conserved in rodents over 30 million years since the mouse-rat lineage divergence.

We conducted RT-qPCR, Northern blotting and RNA-FISH to evaluate expression, transcription variants and subcellular localisation of *Epn* in ESCs. *Epn* showed strong induction within 12 hours of 2i/LIF withdrawal, but its level decreased subsequently (Fig1B-D). In EpiSCs or mouse embryonic fibroblasts (MEFs), *Epn* expression was below the detection limit (Fig1C). Consistent with the UCSC gene annotation, Northern blotting of total ESC RNAs confirmed the expression of a single *Epn* transcript over 1 kb in length (Fig1C, Fig1 – Figure Supplement 1D). Transcription start and end sites of *Epn* mapped by 5’ and 3’ RACE were in agreement with the annotation (Fig1 – Figure Supplement 1E,F). Upon 24 hours post 2i/LIF withdrawal, *Epn* RNA-FISH hybridisation signals displayed predominantly cytoplasmic localisation, but from 48 hours onwards the remaining signals were mostly in the nucleus (Fig1D).

To explore the expression regulation of *Epn*, two inhibitors and LIF were withdrawn singly or dually for 24 hours. In conditions lacking Gsk3 inhibitor CHIRON99021 (CH), *Epn* was upregulated (Fig1E). In LIF/serum, *Epn* expression was twofold higher than in 2i (Fig 1- Figure Supplement 1G) and the addition of CH to LIF/serum culture suppressed *Epn* expression within 24 hours irrespective of the presence of MEK inhibitor PD0325901 (PD) (Fig1 – Figure Supplement 1H). Upon PD/LIF withdrawal, *Epn* expression was maintained for 24 hours then declined (Fig1F). Therefore, *Epn* is suppressed by CH in self-renewing ESCs.

During early mouse development (Boroviak et al., 2015), *Epn* expression peaked at E4.5 and was present in both epiblast and primitive endoderm of the mature blastocyst, and was absent or low in E5.5 post-implantation epiblast (Fig1 – Figure Supplement 2A) and later stages between E7 and E17 (Fig1 – Figure Supplement 2B). Amongst somatic tissues analysed, *Epn* was only detected in kidney, but at a much lower level than in ESCs. We also observed that *Epn* expression is restored upon naïve state resetting from EpiSCs (Guo et al., 2009; Yang et al., 2010) (Fig1 – Figure Supplement 2C,D). We conclude that *Epn* expression is highly specific to ESCs and the early mouse embryo.

LINE and ERVL-MaLR elements are present within the *Epn* proximal promoter region (2kb upstream of TSS) (Fig1- Figure Supplement 1A). Since such repetitive elements gain DNA CpG methylation dramatically during pre-to post-implantation transition (Smith et al., 2014). By examining published data from embryos (Seisenberger et al.; Wang et al., 2014) and ESC progression *in vitro* (Kalkan et al., 2017), we found that CpG methylation gain at the *Epn* promoter was more extensive in the primed E6.5 epiblast (3% to 80%) than the average changes across all promoters (9% to 35%) or the genome (24% to 70%) (Fig 1- Figure Supplement 2E). In contrast, no major CpG methylation gain at *Epn* was present in ESCs 24 hours post 2i withdrawal. These data suggest that *Epn* promoter methylation does not initiate repression, but could contribute to maintain *Epn* silencing in gastrulating epiblast.

### Loss of *Ephemeron* delays exit from naïve pluripotency

Initiation of ESC differentiation in defined media upon withdrawal of self-renewal factors recapitulates features of peri-implantation epiblast development (Kalkan et al., 2017). The naïve state exit latency varies, however, according to the starting self-renewal condition (Dunn et al., 2014; Wray et al., 2011). Higher activity of the core network in PD/LIF compared with 2i results in slower network dissolution, reflecting in later onset of RGd2 downregulation (Dunn et al., 2014). PD/LIF and 2i also feature different levels of *Epn* due to CH mediated suppression in 2i (Fig1E). We therefore generated *Epn* knockout (KO) ESCs via sequential gene targeting (Fig2- Figure Supplement 1) and examined *Epn* KO phenotype for ESCs maintained in each condition. In steady state self-renewal, *Epn* loss did not affect the Rex1-GFP profile in either case (Fig2A,B). Upon transfer to N2B27, *Epn* KO cells displayed delayed downregulation of GFP compared to parental cells, measured at 24h from 2i culture and 40h from PD/LIF culture (Fig2B). At 72 hours, however, GFP expression was fully extinguished from either starting condition (Fig 2 – Figure Supplement 2A). Similarly, delayed GFP downregulation in both culture conditions was also evident upon *Epn* knockdown using siRNAs (Fig 2 – Figure Supplement 2B). To assess the effect of *Epn* depletion functionally, we conducted colony forming assays, in which cells maintained in PD/LIF were subjected to 40 hours culture in N2B27 and then plated at clonal density in 2i/LIF to assay the persistence of ES self-renewal potential (Betschinger et al., 2013). *Epn* KO and knockdown cells both gave rise to substantially more undifferentiated colonies than wild type controls (Fig2C,D, Fig 2 – Figure Supplement 2C). Considered together, these results indicate that the absence of *Epn* impairs timely exit from naïve pluripotency.

**Figure 2:**
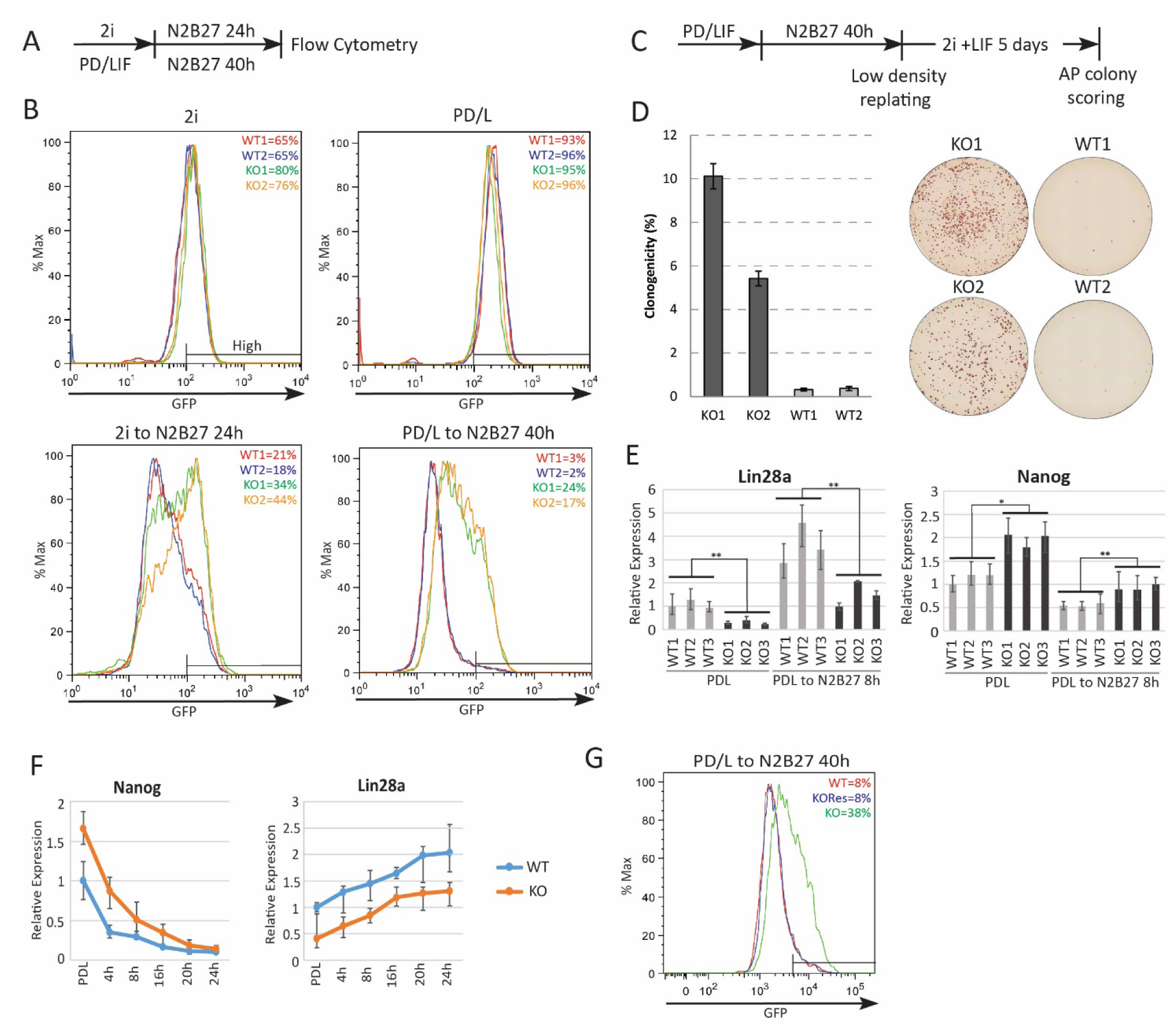
Absence of *Ephemeron* delays exit from naïve pluripotency. A, Experimental scheme for analysing naïve state exit using Rex1GFPd2 reporter cells. B, Rex1-GFP flow cytometry profiles of wild type and *Epn* KO cells in 2i and PD/LIF and during transition from these starting conditions. Two independent clones for wild type and *Epn* KO cells were analysed. Percentage of GFP high cells were quantified. C, Experimental scheme for colony formation assay. D, Colony formation assay for wild type and *Epn* KO cells in 2i/LIF 40 hours post PD/LIF withdrawal. Colonies were stained with alkaline phosphatase (AP), with representative images shown. Percentage clonogenicity was calculated by the number of AP positive colonies divided by the total number of cells plated. Mean+/-SD, n=3. E, *Lin28a* and *Nanog* expression relative to *β-actin* in three independent wild type and *Epn* KO cell lines measured by RT-qPCR. Mean+/-SD, n=3. *p<0.05, **p<0.01, student’s *t*-test. F, *Nanog* and *Lin28a* expression kinetics upon PD/LIF withdrawal in wild type and *Epn* KO cells. Mean+/-SD, n=3. G, Rex1-GFP flow cytometry profile for wild type, *Epn* KO and *Epn* rescue cells 40 hours post PD/LIF withdrawal. Percentage of GFP high cells were quantified.

### Molecular consequences of *Ephemeron* loss

We performed RNA-sequencing and compared the transcriptome of wild type and *Epn* KO ESCs using three independently targeted KO ESC lines and three subclones of the parental wild type ESCs. Twenty-two and Fifty-five genes were significantly differentially expressed between wild type and *Epn* KO cells in PD/LIF and 8 hours after PD/LIF withdrawal respectively (Benjamini-Hochberg adjusted p<0.05, fold change >1.5 or <0.7) (Fig 2 – Figure Supplement 2D). Sixteen of these differentially expressed genes were common to both time points (Fig 2 – Figure Supplement 2E), amongst which, Tcf15 and Lin28a have been associated with exit from the naïve state, with an expression pattern inversely correlated with that of naïve pluripotency factors (Davies et al., 2013; Kumar et al., 2014). *Lin28a* was the most differentially expressed gene among these 16 differentially expressed genes common to both time points, with *Epn* KO cells displaying a twofold reduction in mean expression level (Fig2E). Attenuated downregulation of members of the core naïve transcription factor network is one explanation for delayed exit from ESC state (Kalkan and Smith, 2014). We hypothesised that Lin28a could be a negative regulator of the naïve network. Although Lin28a is commonly considered as a pluripotency factor, its expression is actually increased when cells transiting out of the naïve state *in vivo* and *in vitro* (Boroviak et al., 2015; Kalkan et al., 2017; Marks et al., 2012). We examined expression of naïve pluripotency transcription factors in *Epn* KO cells and found a higher level of *Nanog* mRNA in PD/LIF and 8 hours post withdrawal (Fig2E, Fig 2 – Figure Supplement 2F). To characterise the profile of naïve pluripotency dissolution further in *Epn* KO cells, a PD/LIF withdrawal time course was monitored over 24 hours. The twofold reduction in *Lin28a* mRNA in *Epn* KO cells was constant throughout this time course (Fig2F). Conversely, *Nanog* transcript and protein levels remained higher at 16 hours and 24 hours respectively (Fig2F, see also 3E,F for protein). Mean *Klf2* transcript levels appeared higher in *Epn* KO cells, but it was below statistical significance. Other members of the naïve network showed similar expression profiles in wild type and *Epn* KO cells (Fig 2 – Figure Supplement 2F). Among peri-implantation epiblast markers, upregulation kinetics for *Fgf5* and *Otx2* were unchanged in *Epn* KO cells, but transcripts for *Dnmt3a*, *Dnmt3b* and *Oct6* remained lower in *Epn* KO cells from 16 to 24 hours (Fig 2 – Figure Supplement 2F). Although not statistically significant, *Otx2* transcripts appeared modestly reduced throughout the time course, which could be related to the elevated expression of Nanog (Acampora et al., 2016).

We restored *Epn* expression in KO cells by inserting the *Epn* genomic coding region under the control of human *EF1α* promoter into the deleted locus (Fig 2 – Figure Supplement 3A,B). The rescue cells displayed a wild type exit profile as measured by GFP profile 40 hours post PD/LIF withdrawal (Fig2G). *Lin28a* and *Nanog* expression levels were similar to wild type cells (Fig 2 – Figure Supplement 3C).

To explore the differentiation capacity of *Epn* KO cells, we conducted *in vitro* differentiation assays directing ESCs towards EpiSCs and somatic lineages (Fig2- Figure Supplement 4). Both wild type and *Epn* KO ESCs could be differentiated into EpiSCs using N2B27 supplemented with ActivinA/Fgf2/Xav939 (Sumi et al., 2013) on fibronectin. Such *in vitro* differentiated EpiSCs could be stably propagated over multiple passages, displayed similar morphology and gene expression typical of EpiSCs irrespective of the genotype (Figure 2- Figure Supplement 4A,B). We also applied neuronal, mesendoderm and definitive endoderm differentiation protocols to *Epn* KO ESCs and found that lineage markers were induced, with a slight delay for mesendoderm relative to wild type cells (Figure 2- Figure Supplement 4C-E). Thus delayed downregulation of *Nanog* and *Klf2* observed in all protocols and postponed naïve state exit does not impair subsequent lineage commitment capacity.

### Lin28a and Nanog are in a genetic network with *Ephemeron*

Based on the preceding data, we hypothesised that Lin28a could be a downstream effector of *Epn* and the delayed exit phenotype might be attributed, at least in part, to the elevated starting level and increased perdurance of Nanog. To characterise further the relationship between *Epn*, Lin28a and the naïve transcription factor network, we carried out a series of genetic perturbation experiments and measured both Rex1-GFP reporter dynamics and colony formation upon withdrawal from PD/LIF. Nanog depletion in wild type cells did not substantially alter Rex1-GFP profile, but did reduce the colony formation capacity of cells recovered at 40 hours post PD/LIF withdrawal (Fig3A). Nanog knockdown in *Epn* KO cells partially restored downregulation of Rex1-GFP at 40 hours. Functional ESC exit measured by colony formation capacity 40 hours post PD/LIF withdrawal was fully restored to the reduced level obtained in wild type cells after Nanog knockdown (Fig3A). Knockdowns of other naïve transcription factors, Esrrb, Tfcp2l1 and Klf2, accelerated exit in wild type cells with distinct potencies, and in all cases the phenotype was attenuated in *Epn* KO cells (Fig3- Figure Supplement 1B). Knockdown of Klf4 had no effect in either wild type or *Epn* KO cells starting from PD/LIF. Delayed exit kinetics of Esrrb, Tfcp2l1 and Klf2 knockdown in *Epn* KO cells could be attributed to elevated Nanog. We therefore conducted dual knockdown experiments (Fig3- Figure Supplement 1C). Simultaneous knockdown of Esrrb, Tfcp2l1 or Klf2 together with Nanog largely abolished the effect of *Epn* KO on GFP downregulation (Fig3- Figure Supplement 1C,D). These data are consistent with *Epn* acting via modulation of Nanog expression.

**Figure 3:**
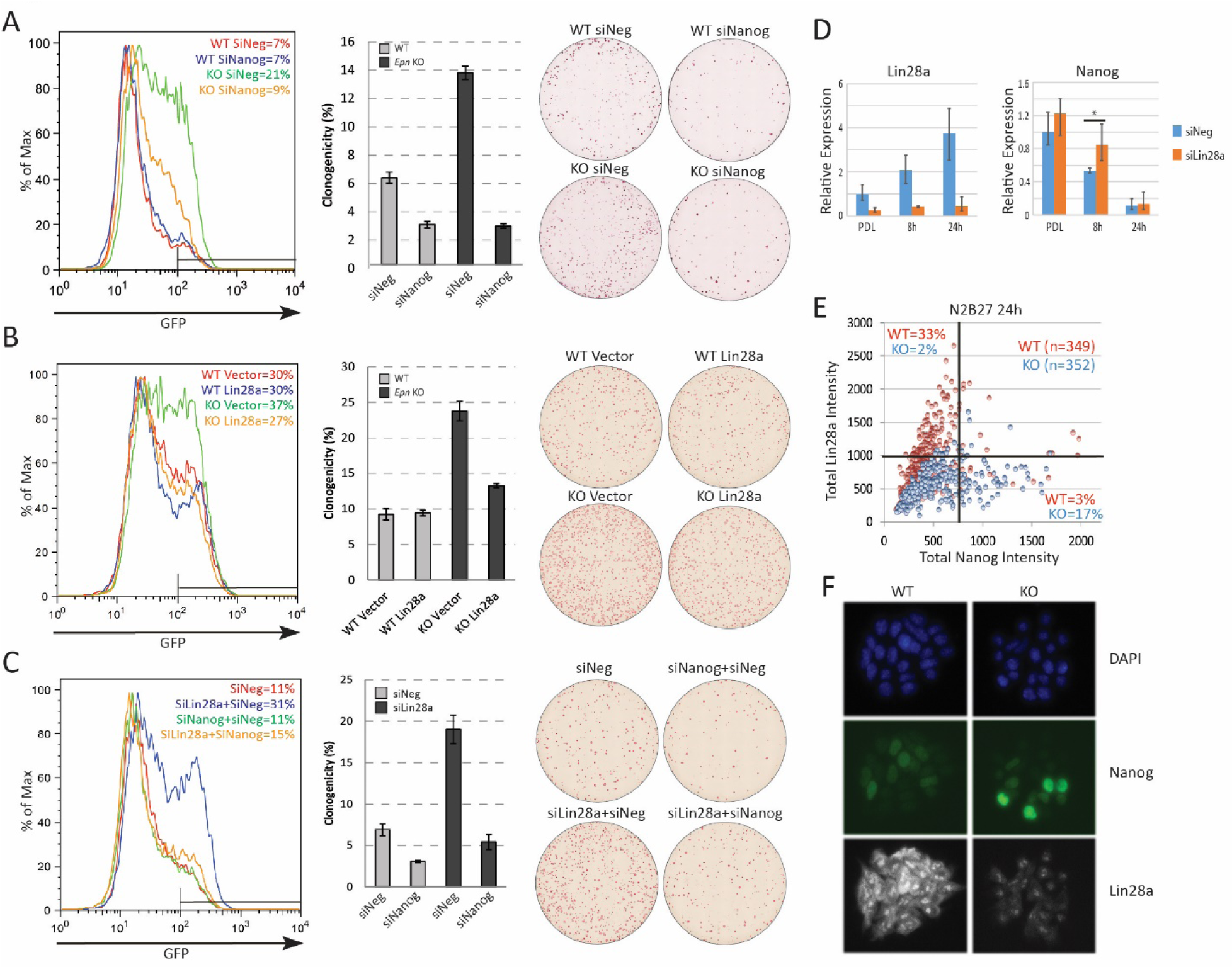
Lin28a is downstream of *Ephemeron* and regulates Nanog expression. A, Rex1-GFP flow cytometry profiles (Left) and colony formation capacity (Right) 40 hours post PD/LIF withdrawal for wild type and *Epn* KO cells transfected with indicated siRNAs. B, Rex1-GFP flow cytometry profiles and colony formation capacity 40 hours post PD/LIF withdrawal for wild type and *Epn* KO cells transfected with Lin28a expression vector. C, Rex1-GFP flow cytometry profile and colony formation capacity 40 hours post PD/LIF withdrawal with Nanog and Lin28a single or dual knockdowns in wild type cells. Quantification of percentage of GFP high cells were shown in A-C. Percentage clonogenicity in A-C is measured by the number of AP positive colonies divided by the total number of cells plated, with representative AP staining images shown. Mean+/-SD, n=3. D, *Lin28a* and *Nanog* expression relative to *β-actin* upon PD/LIF withdrawal in Lin28a knockdown and control cells. Mean+/-SD, n=3. * p<0.05, Student’s *t*-test. E, Correlation of Nanog and Lin28a protein expression immunostaining in wild type and *Epn* KO cells 24 hours post PD/LIF withdrawal. F, Representative images of cells co-immunostained with Nanog and Lin28a and quantified in E.

We investigated whether lowered expression of Lin28a contributes to the slower exit from naïve pluripotency and the increased Nanog expression. We manipulated Lin28a dosage by either overexpression or knockdown in *Epn* KO cells. In wild type cells, Lin28a overexpression had no significant effect. In *Epn* KO cells, however, it restored normal transition kinetics (Fig3B). Conversely, Lin28a knockdown phenocopied *Epn* loss, delaying exit from naïve pluripotency (Fig3C). Concomitant knockdown of Nanog and Lin28a abolished this effect (Fig3C). Lin28a knockdown cells exhibited marginally elevated *Nanog* mRNA in PD/LIF and persistence at higher levels after 8 hours of PD/LIF withdrawal (Fig3D). At the protein level, *Epn* null cultures displayed more cells with high Nanog and low Lin28a expression at the 24-hour time point as quantified by co-immunostaining (Fig3E,F, Fig3- Figure Supplement 1E). Interestingly, Lin28a was detected as concentrated foci in the nucleus and also in the cytoplasm (validated with two independent antibodies), and both nuclear and cytoplasmic expression were induced after PD/LIF withdrawal (Fig3- Figure Supplement 1F). During early embryo development, expression of *Lin28a* and *Epn* are positively correlated, while *Lin28a* and *Nanog* are negatively correlated (Fig3- Figure Supplement 1G)(Ohnishi et al., 2014). These data are consistent with the proposition that Lin28a is genetically downstream of *Epn* and may facilitate exit from naïve pluripotency by accelerating downregulation of Nanog.

To assess whether *Epn* could regulate *Lin28a* or *Nanog* expression directly by promoter localisation (Rinn and Guttman, 2014), we employed chromatin isolation by RNA purification (ChIRP) (Chu et al., 2011). Using this method, we were able to selectively pull down endogenous *Epn* RNA (Fig3- Figure Supplement 2A). However, we did not detect chromatin enrichment at the *Lin28a* or *Nanog* promoter regions (Fig3- Figure Supplement 2B-D). Indeed, no significant enrichment genome-wide was observed in wild type compared to *Epn* KO cells (Fig3- Figure Supplement 2E). Thus we found no evidence that *Epn* functions by chromatin association. However, the H3K4me3 modification was reduced at the *Lin28a* promoter in *Epn* KO cells (Fig3- Figure Supplement 2F).

One explanation for anti-correlated expression could be direct negative regulation of *Lin28a* by Nanog. We therefore inspected two published Nanog chromatin immunoprecipitation (ChIP) sequencing datasets (Chen et al., 2008; Marson et al., 2008) but observed no localisation of Nanog at the *Lin28a* locus (Fig3- Figure Supplement 2G). Furthermore, we did not observe *Lin28a* downregulation in Nanog knockdown cells (Fig 3- Figure Supplement 2H). Therefore, Nanog does not appear to be a direct upstream regulator of Lin28a.

### The function of Lin28a in ESC transition is mediated by suppression of *Mirlet7g*

Lin28a is an RNA binding protein with a well-established function in suppressing maturation of *Mirlet7* family miRNAs (Cho et al., 2012; Viswanathan et al., 2008). We investigated whether the role of Lin28a in naïve state exit is *Mirlet7* dependent. We profiled mature miRNA expression of *Mirlet7* family members using RT-qPCR. Expression of *Mirlet7a, Mirlet7d, Mirlet7e, Mirlet7g* and *Mirlet7i* decreased 24 hours after 2i/LIF withdrawal, coinciding with the increase in Lin28a expression (Fig4A). However, mature miRNA *Mirlet7c* expression was unaffected, suggesting that *Mirlet7c* expression is independent of Lin28a. This observation is in agreement with a recent finding that *Mirlet7c-2*, the major *Mirlet7c* isoform expressed in mouse ESCs, bypasses Lin28a regulation due to lack of a GGAG recognition motif in its loop region (Triboulet et al., 2015). The Lin28a regulated *Mirlet7* miRNAs, but not *Mirlet7c*, are expressed at higher levels in ESCs in 2i/LIF than in LIF/serum (Pandolfini et al., 2016) (Fig4- Figure Supplement 1A), consistent with lower Lin28a in 2i/LIF.

**Figure 4:**
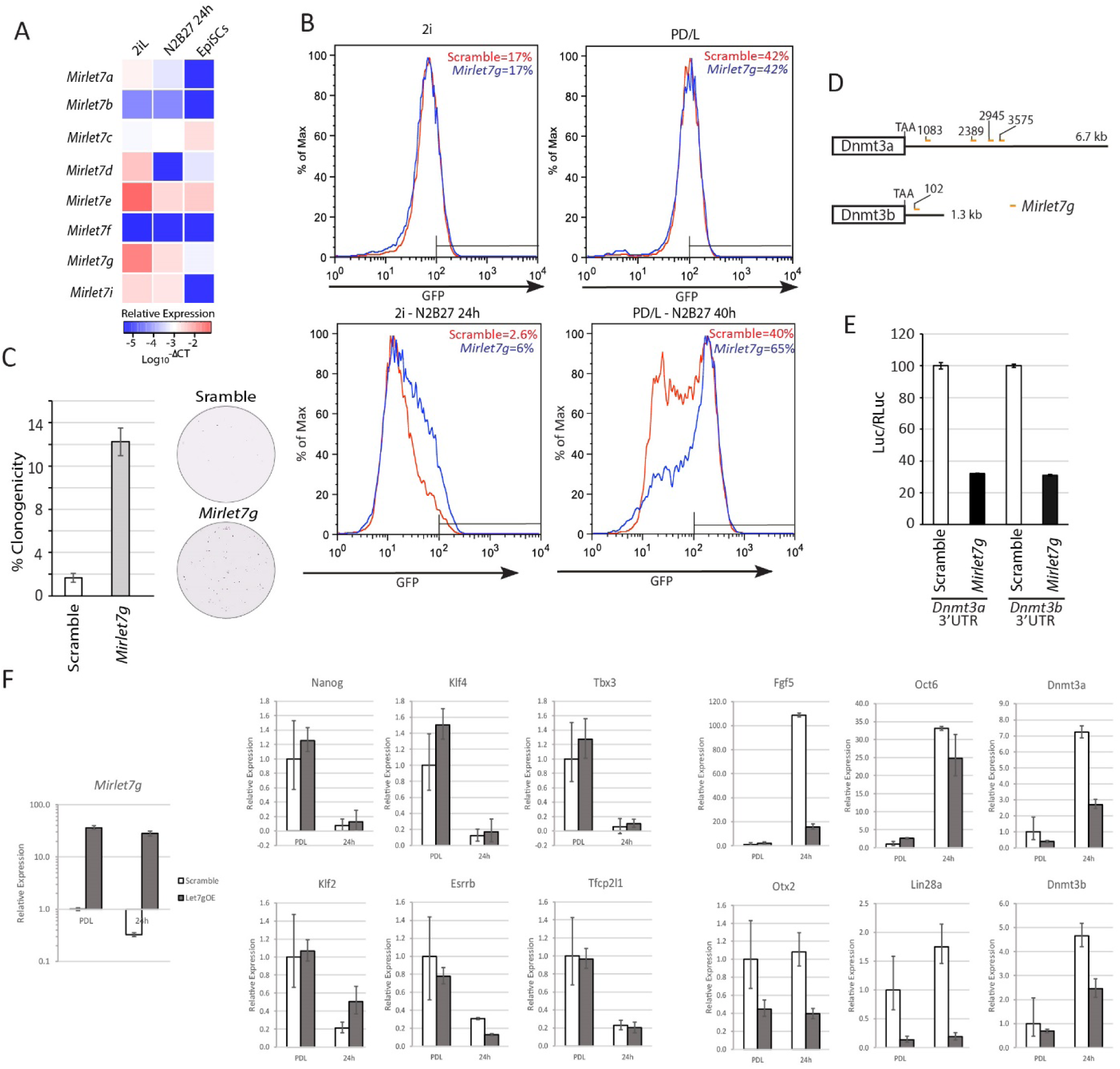
Lin28a function is mediated via members of *Mirlet7* miRNAs. A, Mature *Mirlet7* family microRNA expression quantified by RT-qPCR in 2i/LIF, 24 hours post 2i/LIF withdrawal, and EpiSCs. B, Rex1-GFP flow cytometry profile upon forced expression of mature *Mirlet7g* mimic during transition from 2i and PD/LIF. Quantification of percentage of GFP high cells was shown. C, Colony formation assay in 2i/LIF of cells with forced expression of *Mirlet7g* mimic and control 40 hours post PD/LIF withdrawal. Colonies were stained with alkaline phosphatase (AP). Percentage clonogenicity was calculated by the number of AP positive colonies divided by the total number of cells plated. Mean+/-SD, n=3. D, Predicted target sites of *Mirlet7g* in 3’UTRs of *Dnmt3a* and *Dnmt3b* by RNA22. E, Dual luciferase assay measuring repression by *Mirlet7g* mediated through 3’UTRs of *Dnmt3a* and *Dnmt3b*. Mean+/-SD, n=3. F, Relative expression normalised to *β-actin* of naïve and peri-implantation epiblast associated genes in ESCs with forced expression of *Mirlet7g* mimics. Mean+/-SD, n=3.

To examine the role of Lin28a regulated *Mirlet7* miRNAs in naïve state exit, we transfected ESCs with mature *Mirlet7g* mimic. We used *Mirlet7g* as a representative member since all apart from *Mirlet7e* share the same seed sequence (Fig4- Figure Supplement 1B). Forced expression of *Mirlet7g* in RGd2 cells resulted in delayed GFP downregulation upon both 2i and PD/LIF withdrawal (Fig4B). Elevated ESC colony formation capacity post PD/LIF withdrawal was also observed (Fig4C). To identify the downstream targets of *Mirlet7g*, we searched for genes that are upregulated upon 2i withdrawal in our RNA-sequencing dataset and are either known or predicted *Mirlet7g* targets using the RNA22 tool (Miranda et al., 2006). DNA methyltransferases Dnmt3a and Dnmt3b emerged as prime candidates as has previously been proposed (Kumar et al., 2014). Expression of both increases during transition from both 2i and PD/LIF (Kalkan et al., 2017). *Dnmt3a/3b* transcript levels were lower in *Epn* KO cells than wild type control (Fig2- Figure Supplement 2F). Multiple *Mirlet7g* target sites were predicted by RNA22 within the *Dnmt3a* 3’UTR and one site in the *Dnmt3b* 3’UTR (Fig4D, Fig4- Figure Supplement 1C). ESCs were co-transfected with mature *Mirlet7g* mimic and luciferase constructs containing the entire 3’UTRs of *Dnmt3a* and *Dnmt3b*. *Mirlet7g* reduced luciferase expression by more than 60% relative to the negative control (Fig4E), suggesting that *Dnmt3a/3b* transcripts are indeed *let7-g* targets.

### Dnmt3a and Dnmt3b methylate the Nanog promoter during naïve state exit

Epiblast progression is associated with genome-wide *de novo* methylation during pre-to post-implantation development (Auclair et al., 2014). This phenomenon is recapitulated when naïve ESCs are withdrawn from 2i (Buecker et al., 2014; Kalkan et al., 2017). Previous studies demonstrated hypomethylation of the *Nanog* promoter in mouse ESCs compared to lineage committed cells (Farthing et al., 2008; Yu et al., 2007). We speculated that impeded *de novo* DNA methylation could allow perdurance of *Nanog* expression at the onset of naïve state exit. To investigate this hypothesis, we carried out bisulfite sequencing analysis across the *Nanog* proximal promoter region, 1 kb upstream of the TSS, after siRNA knockdown of Dnmt3a/3b singly or together (Fig5A). We observed a marked reduction of CpG methylation in the -1kb to -761 bp region (region 1) of the *Nanog* proximal promoter 40 hours after PD/LIF withdrawal (Fig5A). At this time point, control cells exhibited 40% CpG methylation at scored sites, whereas Dnmt3b depleted cells displayed 18% CpG methylation and Dnmt3a or Dnmt3a/b double knockdown cells showed only around 8%. Effects were less obvious in the -538 bp to +18 bp region (region 2), which was barely methylated at this time point. *Epn* KO cells also exhibited reduced methylation, with region 1 again showing a more prominent reduction (Fig5 – Figure Supplement 1A). These data suggest that Dnmt3a and Dnmt3b have overlapping roles in mediating *de novo* methylation at the *Nanog* proximal promoter.

**Figure 5:**
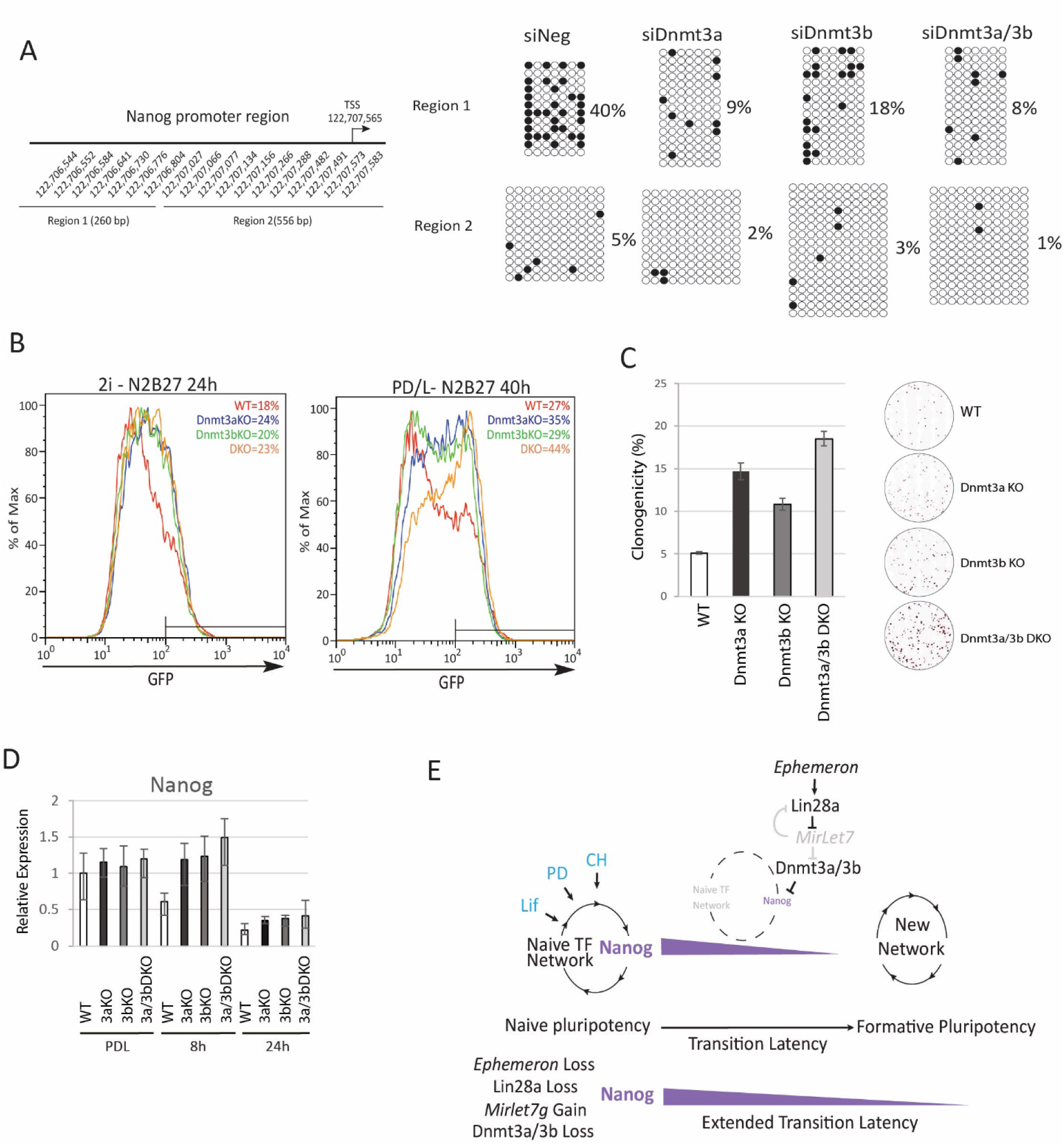
Loss of Dnmt3a and Dnmt3b delays naïve state exit associated with transient persistence of Nanog expression. A, Bisulfite sequencing analysis of *Nanog* proximal promoter CpG island DNA methylation in Dnmt3a and Dnm3b single and dual knockdown cells at 40 hours post PD/LIF withdrawal. The positions of cytosines analysed (mm10) are indicated on the left panel. Black and while circles represent methylated and unmethylated cytosine respectively. B, Rex1-GFP2 flow cytometry profiles of *Dnmt3a* and *Dnmt3b* single and dual KO cells withdrawn from 2i or PD/LIF for 24 and 40 hours respectively. Percentage of GFP high cells were quantified. C, colony formation capacity 40 hours post PD/LIF withdrawal for *Dnmt3a* and *Dnmt3b* single and compound KO cells. Percentage clonogenicity was measured by the number of AP positive colonies formed divided by the total number of cells plated, with representative AP staining images shown. Mean+/-SD, n=3. D, Expression of *Nanog* relative to *β-actin* in *Dnmt3a* and *Dnmt3b* single and compound KO cells quantified by RT-qPCR. Mean+/-SD, n=2. E, schematic representation of the inferred *Epn* pathway.

To explore the role of *de novo* DNA methylation in ESC transition, we investigated functional consequences of *Dnmt3a* and *Dnmt3b* depletion. We generated *Dnmt3a* and *Dnmt3b* single and compound knockouts in RGd2 ESCs using CRISPR/Cas9. Using two guide RNAs (gRNAs), we generated deletions of highly conserved PC and ENV motifs (motifs IV and V) within the catalytic domain for both Dnmt3a and Dnmt3b, recapitulating the previously characterised *Dnmt3b* and *Dnmt3b* mutant gene structures (Okano et al., 1999) (Fig5 – Figure Supplement 1B). *Dnmt3a* and *Dnmt3b* single and double KO cells exhibited delayed GFP downregulation (Fig5B). Colony formation capacity of the single and double KO cells 40 hours post PD/LIF withdrawal confirmed a delay in the extinction of ESC identity (Fig5C). Interestingly, however, the delayed GFP downregulation did not endure (Fig5 – Figure Supplement 1C). Absence of Dnmt3a and Dnmt3b singly or together was associated with increased *Nanog* expression (Fig5D). At 8 hours after PD/LIF withdrawal, *Nanog* mRNA in *Dnmt3a* and/or *Dnmt3b* mutants was equivalent to wild type cells in PD/LIF, whereas wild type cells had downregulated *Nanog* expression by 50% (Fig5D). We also observed elevated expression of *Tfcp2l1*, *Klf2*, *Klf4* and *Tbx3* in *Dnmt3a/3b* single or compound KO cells (Fig5 – Figure Supplement 1E). The promoters of these genes are methylation refractory in 2i withdrawal time course (Kalkan et al., 2017). Therefore, the elevated expression should be secondary to some other factor(s) such as increased Nanog expression. *Dnmt3a/3b* compound KO also resulted in impeded upregulation of peri-implantation markers such as *Fgf5*, *Oct6* and *Otx2* at 24 hours post PD/LIF withdrawal (Fig5 – Figure Supplement 1F). These data indicate that *de novo* DNA methylation facilitates timely progression from the ESC state. Importantly, however, *de novo* methylation by Dnmt3a/3b is not essential for the exit from naïve pluripotency.

## Discussion

In this study, we identified a genetic network that connects a lncRNA, *Ephemeron*, with known players in post-transcriptional and epigenetic regulation. *Epn* sits at the apex of this cascade, upstream of Lin28a/*Mirlet7g* and Dnmt3a/3b (Fig5E), and ultimately contributing to downregulation of the potent naïve transcription factor Nanog. *Epn* facilitates upregulation of Lin28a, although how this is achieved remains unclear. Lin28a in turn suppresses sub-family members of the *Mirlet7* miRNAs such as *Mirlet7g*, targets of which include *de novo* DNA methyltransferases Dnmt3a and Dnm3b. Increased Dnmt3a/3b activity correlates with *Nanog* proximal promoter CpG methylation, which may consolidate the lowered expression trigged by withdrawal of 2i/LIF. This mechanism provides an additional dimension to the multi-layered molecular machinery that expedites the irreversible ESC transition from naïve to formative pluripotency (Jang et al., 2017; Kalkan et al., 2017; Kalkan and Smith, 2014).

ES cell maintenance is robust due to parallel pathway wiring (Dunn et al., 2014). Progression from such a resilient state requires a powerful and coordinated dissolution machinery. Our findings indicate that one component is the *Epn* cascade. Gsk3 inhibition by CH represses *Epn* and thereby Lin28a, consistent with insulation of the naïve transcription factor network (Wray et al., 2011). In contrast, ESCs cultured in PD/LIF express *Epn* and Lin28a but without overt consequence, presumably due to the potent self-renewal environment of Stat3 activation and MEK inhibition that sustain expression of Nanog and other naïve factors. However, loss of *Epn* in PD/LIF resulted in elevated Nanog.

LncRNAs are more tolerant to TE integration than protein coding genes and TE could drive the more rapid evolution than in protein coding genes (Kelley and Rinn, 2012; Necsulea et al., 2014). *Epn* is comprised of 76.4% TEs, compared to the average of 41.4% TE composition in the mouse genome and of 33% reported for mouse multi-exon lincRNA sequences (Kelley and Rinn, 2012). The aligned sequence between *Epn* and the rat transcript from the syntenic region includes ERVK LTR and SINE B2 elements. These sequences have been preserved for over 30 million years since mouse-rat lineage divergence, which could be indicative of functional constraint on *Epn* sequence and these domesticated TEs. Interestingly, non-coding transcripts harbouring TE sequences are enriched in ESCs and early embryo development for both mouse and human (Fort et al., 2014; Göke et al., 2015; Kelley and Rinn, 2012) and in several instances have been proposed to regulate pluripotency through different mechanisms (Durruthy-Durruthy et al., 2016; Fort et al., 2014; Lu et al., 2014).

However, *Epn* is only found in rodents. In fact, species-specific lncRNAs comprise the majority of the lncRNAs discovered, in particular within the primate branches (Necsulea et al., 2014). Due to their more rapidly evolving nature, it is thought that lncRNAs are more likely to acquire species-specific and lineage-restricted functions and several such examples have recently been characterised (Durruthy-Durruthy et al., 2016; Paralkar et al., 2014; Rani et al., 2016). The rodent specificity of *Epn* might be related to the more rapid progression from pre-implantation epiblast to gastrulation in rodents and the associated requirements for acute extinction of the naïve pluripotency programme (Smith, 2017).

In common with Lin28a (Shinoda et al., 2013), *Epn* is dispensable for development *in vivo*, as we obtained a Mendelian ratio of homozygous *Epn* mutant mice from heterozygous intercrosses (5:18:7, number of wild type: heterozygous: homozygous offspring). Thus ESCs provide a sensitised platform for dissecting redundant individual elements within a multi-layered control machinery for pluripotency regulation (Leeb et al., 2014; Martello and Smith, 2014).

Lin28a is known as a human somatic cell reprogramming factor that acts to suppress *Mirlet7* miRNA family members, which are highly expressed in differentiated cells (Melton et al., 2010; Yu et al., 2007). However, *Lin28a* is expressed at a low level in ground state mouse ESCs (Marks et al., 2012) and pre-implantation epiblast, but at high levels in post-implantation epiblast and EpiSCs (Boroviak et al., 2015). The expression pattern is consistent with our evidence that up-regulation of Lin28a at the onset of mouse ESC differentiation functions to facilitate transition from the naïve state. During human iPSC generation, it is plausible that Lin28a promotes acquisition of primed pluripotency, the endpoint for current human somatic cell reprogramming.

Lin28a itself is a target of *Mirlet7* miRNAs (Kumar et al., 2014; Melton et al., 2010), and this double-negative feedback loop can act as a bimodal switch to facilitate network transition. Our findings are consistent with the recent report that loss of Lin28a reduces ESC heterogeneity in serum/LIF, an effect is mediated by *Mirlet7g* (Kumar et al., 2014). However, independent of *Mirlet7*, Lin28a can post-transcriptionally regulate the expression and/or translation of many RNAs (Cho et al., 2012; Zhang et al., 2016), that could also contribute to naïve state exit.

We observed that *de novo* DNA methyltransferases Dnmt3a/3b are targets of *Mirlet7g*. Naïve ESCs (Ficz et al., 2011; Habibi et al., 2013; Leitch et al., 2013) and pre-implantation epiblast (Monk et al., 1987; Sanford et al., 1987) display global DNA hypomethylation. *De novo* methyltransferases Dnmt3a/3b are lowly expressed and dispensable in ESCs (Okano et al., 1999). However, the post-implantation epiblast rapidly acquires global DNA methylation and this process is dependent on Dnmt3a/3b (Auclair et al., 2014). A similar trend is also observed upon naïve ESC withdrawal from 2i (Kalkan et al., 2017). Loss of Dnmt3a and Dnmt3b singly and in combination delay naïve state exit. The role of *de novo* methylation in facilitating the ESC state exit may be exerted on specific naïve pluripotency associated factors, such as Nanog. It is noteworthy, however, that this effect is transient and ESC identity does not persist. Therefore, although *de novo* DNA methylation facilitates the rapid dissolution of ESC identity, consistent with placement of Dnmt3a/3b downstream of *Epn*, it is not required for the exit from naïve pluripotency.

In summary, we have mapped a genetic interaction pathway consisting of a novel lncRNA, proteins and miRNAs serving as an integral part of the multi-layered molecular machinery that propels mouse ESCs towards lineage competence. The defined mouse ESC system for phased progression of pluripotency is a sensitive experimental platform for the functional annotation of lncRNAs. We speculate that the fine-tuning effect of *Epn* may be representative of lncRNA actions on specific cellular processes.

## Experimental Procedures

### Targeting, expression and gRNA vector construction

BAC RP24-353A19 (C57BL/6J background) was obtained from Wellcome Trust Sanger Institute for constructing the *Epn* knockout targeting vectors by recombineering using bacterial strain EL350 (Lee et al., 2001). Floxed drug resistant cassettes containing hygromycin B phosphotransferase gene (Hygro) or Blasticidin S-resistance gene (Bsd) were PCR amplified using chimeric primers miniU and miniD (Supplementary file 1A) harbouring 80 bp mini-homologies to the genomic region flanking *Epn* locus. The purified PCR fragments *loxP*-PGK-Hygro-bghpA-*loxP* and *loxP*-PGK-Bsd-bghpA-loxP were used to replace the entire genomic region of *Epn* locus with the drug resistant cassettes. The retrieval homology arms were PCR amplified using primers ReUF and ReUR for upstream and ReDF and ReDF for downstream mini-arms (Supplementary file 1B). The mini arms were subsequently cloned into pBS-MC1-DTA vector by 3-way ligation using restriction enzymes, SpeI, HindIII and XhoI. The mini-arm containing vector was linearised by HindIII and used to retrieve the targeting vectors from the modified BACs, giving rise to the final targeting vectors, HygroTV and BsdTV.

Lin28a overexpression vector was constructed by PCR cloning mouse *Lin28a* from cDNA using forward primer AATTGTCGACATGGGCTCGGTGTCCAACCAGCAGT and reverse primer AATTGCGGCCGCTCAATTCTGGGCTTCTGGGAGCAG and cloned into pENTR2B vector. It was subsequently cloned into *PiggyBac*-based expression vector using Gateway LR clonase (Thermo Fisher 11791020) to generate pCAG-Lin28a-pA:PGK-hygro-pA plasmid.

The gRNA design was conducted using online CRISPR gRNA design tool https://www.dna20.com/eCommerce/cas9/input. The chosen gRNAs were based on the minimal off-target scores. For both Dnmt3a and Dnmt3b, the deletion was designed to recapitulate the original Dnmt3a and Dnmt3b KO ES cells (Okano et al., 1999), with the highly conserved PC and ENV motifs (motifs IV and V) within the catalytic domain deleted. The gRNAs were generated by annealing the indicated oligos (Supplementary file 2A), which were subsequently ligated into pX458 vector (Addgene) digested with BbsI. The constructs were sequence validated before transfection.

### Cell Culture

ESCs were cultured on 0.1% gelatin in 2i/LIF medium (NDiff B27 base medium, Stem Cell Sciences, cat. SCS-SF-NB-02, supplemented with 1 μM PD0325901, 3 μM CHIR99021, and 20 ng/ml LIF) as described (Ying et al., 2008). For gene targeting, ESC were maintained with serum containing medium supplemented with 2i/LIF as before (KO-DMEM high glucose, 15% FCS, 2 mM L-Glutamine, NEAA, 1mM Sodium Pyruvate (Life Technologies), 100mM β-Mercaptoethanol (Sigma). Correctly targeted clones were transferred to N2B27 based 2i/LIF medium for expansion and experimentation. The RGd2 reporter wild type subclones and *Epn* KO ESC clones are of V6.5 origin (RRID:CVCL_C865). An independent wild type RGd2 reporter line is of E14 origin (RRID:CVCL_C320). All cell lines are mycoplasma negative and will undergo STR profiling.

### Naïve pluripotency exit assays

ESCs were plated at 1 × 10^4^/cm^2^ in 2i/LIF or PD/LIF. The next day, cells were carefully washed with PBS before switching to NDiff B27 medium. Rex1-GFP profile was analysed at indicated time point in at least two independent experiments on a Cyan or Fortessa FACs analyser and the GFP high population was quantified and indicated in all flow cytometry profiles. Live dead discrimination was performed using Topro-3. For clonal assay, post 24h or 40h 2i or PD/LIF withdrawal respectively, 300-500 cells were plated per well of a 12 well plate coated in Laminin (1:100 dilution, Sigma L2020) and cultured in 2i/LIF for 6 days. Alkaline Phosphatase staining (Sigma 86R-1KT) was conducted to detect ES colonies. AP-stained plates were imaged using an Olympus IX51, DP72 camera with CellSens software and subsequent colony counting was conducted manually using ImageJ software.

### EpiSC derivation from ESCs and EpiSC resetting

ESCs were plated at 1×10^4^/cm^2^ in 2i/LIF on gelatin coated plate. The next day, cells were washed with PBS and before medium switch to NDiff B27 medium supplemented with 20ng/ml Activin A and 12ng/ml Fgf2 together with 2μM XAV939 (Sigma, X3004), A/F/X. Cells were then passaged to fibronectin coated plate in A/F/X medium. EpiSCs were passaged for at least seven times before gene expression analysis and resetting. For EpiSC resetting, EpiSCs were stably transfected with GY118F construct by *piggyBac* transposition (Yang et al., 2010). 1 × 10^4^ cells were plated in a one well of a 12 well plate in A/F/X, the next day, 2i plus GCSF was supplied to initiate resetting.

### Differentiation assays

For neuronal differentiation, ESCs were plated at 1 × 10^4^/cm^2^ in NDiff B27 medium on laminin (1:100 PBS) for up to 4 days for gene expression analysis. For mesendoderm differentiation, cells were plated at 0.6x10^4^/cm^2^ in NDiff B27 based medium containing 10ng/ml ActivinA, 3μM CHIR99021 on Fibronectin for up to 4 days for gene expression analysis. For definitive endoderm differentiation, cells were plated at 1.5x10^4^/cm^2^ in NDiff B27 based medium containing 20ng/ml ActivinA, 3μM CHIR99021, 10ng/ml FGF4, 1μg/ml Heparin, 100nM PI103. On day 2, the media was switched to SF5 based medium containing 20ng/ml ActivinA, 3μM CHIR99021, 10ng/ml FGF4, 1μg/ml Heparin, 100nM PI103 and 20ng/ml EGF2. Per 100ml SP5 basal medium, it contains 500μl N2, 1ml B27 without VitamineA supplement, 1% BSA, 1ml L-glutamine and 100μl β-mercaptoethanol. Detailed protocols can be found in Mulas et al (Mulas et al., 2016).

### SiRNA, miRNA mimics and plasmid transfection

SiRNAs and miRNA mimics were obtained from Qiagen and the catalogue numbers are listed in Supplementary file 3. Transfection was performed using Dharmafect 1 (Dharmacon, T-2001-01) in a reverse transfection protocol with the final siRNA or miRNA mimics concentration to be 10nM. Two siRNA combination were used per transfection for each target gene knockdown.

Plasmid transfection was performed using Lipofectamine 2000 (Life Technologies) following the manufacturers protocol. For *piggyBac* based stable integration, a *piggyBac* transposon and hyperactive PBase (hyPBase) ratio of 3:1 was used.

### Generation of Dnmt3a and Dnmt3b KO ESCs with CRISPR/Cas9

A pair of gRNA containing plasmids based on px458 designed were transfected using Fugene HD (Promega). 100 ng of each plasmid were transfected with 0.6 ul Fugene HD (1:3 ratio) to 2x10^5^ ESCs in suspension in 2iL medium overnight. The next day, the media was refreshed and 48 hours post transfection, 1,000 GFP high cells were sorted into a well of a six well plate for colony formation. Individual colonies were picked and genotyping was conducted from extracted genomic DNA by triple primer PCR to identify clones with designed deletion (Supplementary file 2B). For Dnmt3a KO, deletion resulted in genotyping PCR product to shift from 760bp representing the wild type allele to 1132 bp. For Dnmt3b KO, deletion resulted in genotyping PCR product to shift from 344bp representing the wild type allele to 653bp. Only homozygous mutants were chosen for subsequent experimentation.

### Southern blotting

Genomic DNA of individually picked ESC clones was extracted and digested with *Xmn*I, size-fractionated on a 0.8% agarose gel and transferred to Hybond-XL blotting membrane (Amersham) using standard alkaline transfer methods. The 5’ and 3’ external probes were generated by PCR with primer sequences shown in Supplementary file 4A. Southern blot hybridization was conducted as described previously (Li et al., 2011).

### Northern blotting

10 μg of purified RNA was resolved by denaturing formaldehyde agarose gel electrophoresis with MOPS buffer. RNA was transferred to Hybond-XL (GE Healthcare, RPN2020S) membrane in 1xSSC buffer overnight by capillary transfer. RNA was UV cross-linked to the membrane and pre-hybridised with Expresshyb (CloneTech, 636831) for 2 hours at 65°C. The DNA probe was generated by PCR (primers are shown in Supplementary file 4B) and 25ng of probe DNA was labelled with [^32^P]-dCTP using Radprime DNA labeling system (Invitrogen, 18428-011). The free-nucleotide was removed from labelled probe using G-50 column (GE Healthcare,27-5330-01), and was heat-denatured followed by snap cooling. The probe was added to the pre-hybridised membrane and incubated overnight at 65°C in a rolling incubator. Membrane was washed with wash buffer containing 0.1 x SSC and 0.1% SDS 3 times at 65°C with 10 min intervals. The membrane was placed in a phosphoimager and exposed for at least overnight at -80°C before scanned using Typhoon 9410 phosphoimager system (GE Healthcare).

### 5’ and 3’ RACE

5’ RACE was conducted using 5'-Full RACE Core Set (Takara, #6122) following manufacture’s protocol. The sequences for RT-primer and nested PCR primers A1, A2, S1, and S2 are shown in Supplementary file 5A. 3’ RACE was conducted by using a polyT RT-primer with a unique sequence tag to synthesis cDNA. The 3’ end region was PCR amplified using a primer specific to the RT-primer and a gene specific primer. The primers are shown in Supplementary file 5B. Both 5’ and 3’ RACE PCR products were cloned into plasmids using Zero blunt TOPO PCR cloning kit (Life Technologies, 451245) for subsequent sequencing.

### RNA extraction, reverse transcription and Real-time PCR

Total RNA was isolated using Trizol (Invitrogen) or RNeasy kit (Qiagen) and DNase treatment was conducted either after RNA purification or during column purification. cDNA was transcribed from 0.5˜1 ug RNA using SuperScriptIII (Invitrogen) and oligo-dT priming. Real-time PCR was performed using StepOnePlus machine (Applied Biosystems) with Fast Sybrgreen master mix (Applied Biosystems). Target gene primer sequences are shown in Supplementary file 6. Expression level were normalised to Actinβ. Technical replicates for at least two independent experiments were conducted. The results were shown as mean and standard deviation calculated by StepOnePlus software (Applied Biosystems). The cDNA library for E7-E17 embryos and adult somatic tissues were purchased from Clontech (Mouse Total RNA Master Panel, cat. no. 636644).

### RNA-FISH

RNA-FISH was conducted using ViewRNA ISH Cell Assay for Fluorescence RNA *In Situ* Hybridization system (Affymetrix Panomics) with modifications and imaged on a DeltaVision Core system (Applied Precision), as described in Bergmann *et al*., 2015. The probe set used for *Ephemeron* was VX1-99999-01.

### Mature miRNA expression profiling

Total RNA was extracted using Trizol (Thermo Fisher 15596026). 1 ug RNA was reverse transcribed using Taqman^®^ MicroRNA Reverse Transcription Kit (Thermo Fisher 366596). Mature miRNA expression was analysed using Taqman® Array Rodent MicroRNA A+B Cared Set V3.0 (Thermo Fisher 444909).

### Luciferase assay

The Entire 3’UTR of both *Dnmt3a* and *Dnmt3b* were PCR cloned downstream of the firefly luciferase coding region into pGL3 vector. For *Dnmt3a* 3’UTR, forward primer AATTGGCCGGCCGGGACATGGGGGCAAACTGAAGTAG and reverse primer AATTGGATCCGGGAAGCCAAAACATAAAGATGTTTATTGAAGCTC were used for PCR cloning. For *Dnmt3b* 3’UTR, forward primer AATTGGCCGGCCTTCTACCCAGGACTGGGGAGCTCTC and reverse primer AATTGGATCCTTATAGAGAAATACAACTTTAATCAACCAGAAAGG were used for PCR cloning. The pGL3 vector without the 3’UTR clone was used as a control. Each firefly luciferase construct (500ng) together with Renilla luciferase construct (10ng) were con-transfected with either *Mirlet7g* mimic or scramble control (20nM). The firefly and Renilla luciferase activity was determined by dual luciferase assay (Promega, catalogue no. E1960) 48h post-transfection.

### Immunostaining

Cells were fixed in 4% paraformaldehyde for 10 min at room temperature and were blocked with blocking buffer (5% semi-skimmed milk with 0.1% Triton in PBS) for 2 hours at room temperature. Primary antibodies were diluted in blocking buffer and incubated at 4°C overnight. Primary antibody was carefully washed away with 0.1% Triton in PBS three times with 10 min incubation between each wash. Secondary antibody diluted in blocking buffer (1:1000) was incubated at room temperature for 1 hour followed by 3 washes with 0.1% Triton in PBS. Nuclei were counterstained with DAPI. Primary antibodies used were Nanog (eBioscience, 14-5761, RRID:AB_763613, 1:200) and Lin28a (Cell signalling, 3978, RRID:AB_2297060, 1:800; 8706, RRID:AB_10896850, 1:200). Images from random fields were taken with Leica DMI3000 and the images from different fields at each time point were combined and analysed using CellProfiler software (Broad Institute, RRID:SCR_007358) to conduct nuclear and cytoplasmic compartmentalisation and total fluorescent intensity for each sub-cellular compartments as well as the whole cell for each cell was extracted for correlation analysis.

### Chromatin isolation by RNA purification (ChIRP)

The antisense oligo probes were selected with GC content in the range of 40%-50% in regions of the *Epn* exons without repetitive sequences (Figure 1- Figure Supplement 1A). The probes sequences are in shown in Supplementary file 7. CHIRP was conducted following published protocol (Chu et al., 2011). The data is available at the NCBI Gene Expression Omnibus (accession number: GSE90574). The link to the data is as follows: https://www.ncbi.nlm.nih.gov/geo/query/acc.cgi?token=cralcwwqvtaptgv&acc=GSE90574

### ChIP

The experimental procedure was conducted as described previously (Betschinger et al., 2013). 2 ug of H3K4me3 antibody (Diagenode pAb-003-050) and IgG control (Santa Cruz, sc-2345) was used for 4×10^6^ cells per ChIP. qPCR was performed with primers shown in Supplementary file 8.

### *Nanog* promoter DNA methylation analysis

Genomic DNA was extracted using GenElute Mammalian Genomic DNA miniprep kit (Sigma G1N70-1KT). 500ng purified genomic DNA was treated with sodium bisulfite to convert all unmethylated cytosine residues into uracil residues using Imprint DNA modification Kit (Sigma, MOD50-1KT) according to the manufacturer's protocol. *Nanog* proximal promoter regions (Region 1 and 2 as indicated in Figure 5a) were amplified using a nested PCR approach with KAPA HiFi Uracil+ Readymix (KapaBiosystems, KK2801). The PCR condition for both nested rounds of PCR is as follows: denaturation at 98°C for 5 minutes followed by 10 cycles of gradient PCR, 98°C for 15 seconds, 62°C (starting annealing temperature) for 15 seconds with annealing temperature reduced by 1°C per cycle and 72°C for 1.5 minutes. Followed by this, a 35 cycles of 98°C for 15 seconds, 58°C for 15 seconds and 72°C for 1.5 minutes were conducted. 2μl first round PCR product was used as template for the nested PCR. All primer sequences are shown in Supplementary file 9. The PCR products were verified and purified by gel electrophoresis and subsequently subcloned into PCR4.1 TOPO vector (Invitrogen) according to the manufacturer's protocol. Reconstructed plasmids were purified and individual clones were sequenced (Eurofins).

### Transcriptome sequencing and analysis

Total RNA was isolated with RNeasy RNA purification kit (Qiagen). Ribo-zero rRNA depleted RNA was used to generate sequencing libraries for wild type and Ephemeron knockout cells in PD/LIF and 8 hours withdrawal from PDL from three independent cell lines. Single end sequencing was performed and the reads were mapped using NCBI38/mm10 with Ensembl version 75 annotations. RNA-seq reads were aligned to the reference genome using tophat2. Only uniquely mapped reads were used for further analysis. Gene counts from SAM files were obtained using htseq-count with mode intersection non-empty, -s reverse. Differential gene expression analysis was conducted using Bioconductor R package DESeq2 version 1.4.5. DESeq2 provides two P-values, a raw P-value and a Benjamini-Hochberg P-value (adjusted p value). An adjusted p-Value threshold of 0.05 was used to determine differential gene expression (95% of the results are not false discoveries, error rate 0.05 =5%). The data is available at the NCBI Gene Expression Omnibus (accession number: GSE90574). https://www.ncbi.nlm.nih.gov/geo/query/acc.cgi?token=cralcwwqvtaptgv&acc=GSE90574

### *Epn* promoter CpG methylation analysis

Using published genome-wide bisulpite sequencing data (Kalkan et al., 2017; Seisenberger et al.; Wang et al., 2014), *Epn* promoter region was defined as the 2kb region upstream of the TSS and the percentage of CpG methylation within the region was quantified. For promoter average, percentage of CpG methylation around the 2kb promoter region of each annotated gene was quantified and averaged for all values. For genome average, percentage of CpG methylation of all 50kb tiling windows was quantified and averaged all values.

## Acknowledgments

We thank Kosuke Yusa and Graziano Martello for comments on the manuscript. We are grateful to Carla Mulas for assisting the miRNA expression plot, Yiping Zhang for lncRNA candidate prediction analysis and Rosalind Drummond for technical support. We thank Heather Lee for providing Dnm3ta and Dnmt3b siRNAs. We also thank Peter Humphreys and Andy Riddell for technical support for imaging analysis and flow cytometry respectively. We thank Nicholas Ingolia for useful discussion on ribosomal footprinting on Epn. A.S is supported by Medical Research Council (G1100526/1), Biotechnology and Biological Sciences Research Council (BB/M004023/1), European Commission (HEALTH-F4-2007-200720 EUROSYSTEM), and Wellcome Trust (091484/Z/10/Z). L.H is supported by National Cancer Institute (R01 CA139067, 1R21CA175560-01) and California Institute of Regenerative Medicine (RN2-00923-1), American Cancer Society (123339-RSG-12-265- 01-RMC), Tobacco-related Disease Research Program (21RT-0133). D.L.S. is supported by NIGMS 42694 and NCI 5PO1CA013106-Project 3. T.K. is supported by programme grants from Cancer Research UK (C6/A18796) and European Research Council CRIPTON Grant (268569) and supported by core grants from the Wellcome Trust (092096) and Cancer Research UK (C6946/A14492). The Cambridge Stem Cell Institute receives core funding from the Wellcome Trust and the Medical Research Council. M.A.L. was a Siebel postdoctoral fellow at the University of California, Berkeley and a Sir Henry Wellcome postdoctoral fellow (096125/Z/11/Z). A.S. is a Medical Research Council Professor.

---

**Fig1 – Figure Supplement 1:**
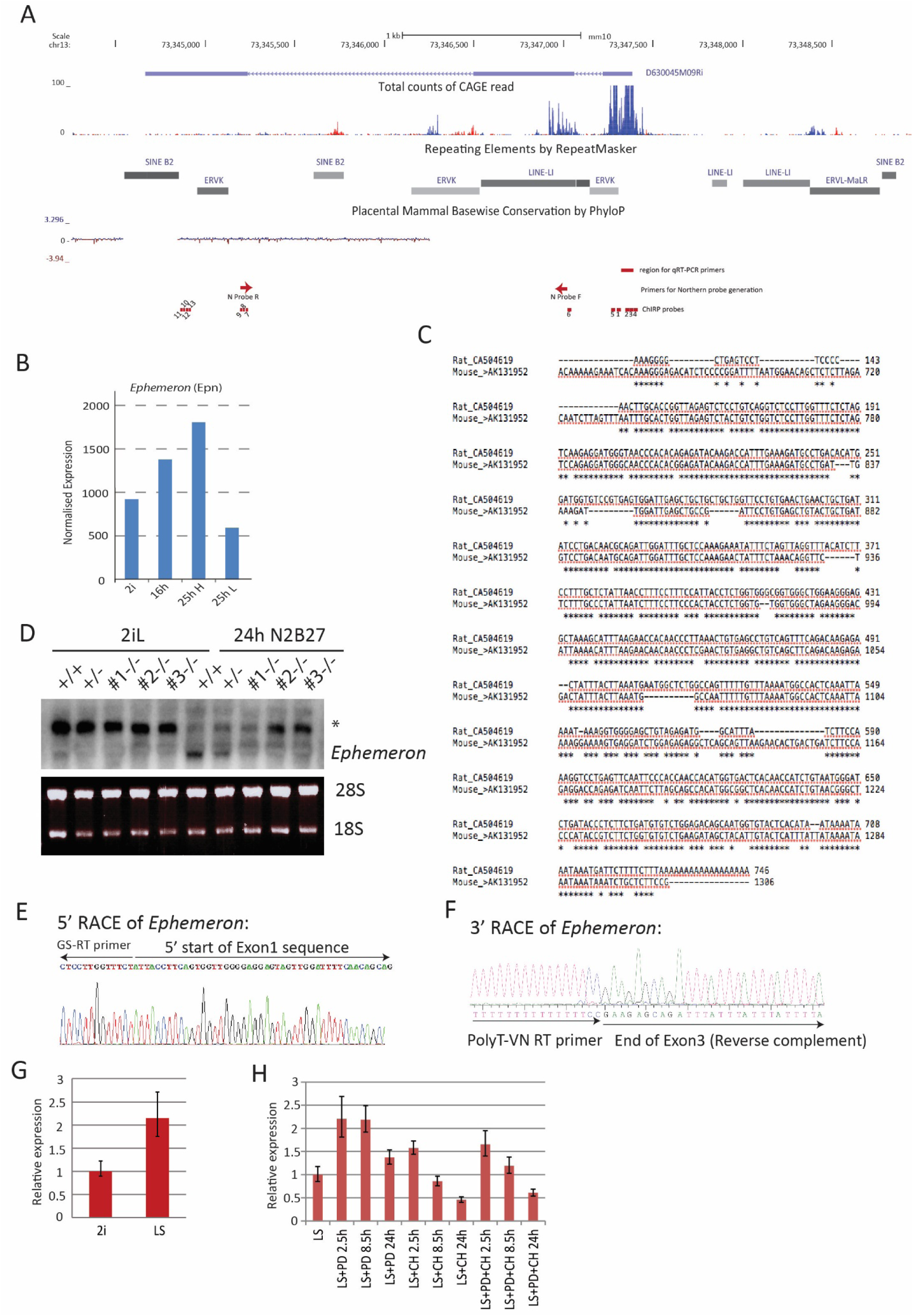
Molecular characterisation of *Ephemeron*. A, *Epn* locus structure, TE content and mammalian conservation. The positions of Northern blotting, RNA-FISH and ChIRP probes were indicated. Note that Northern blotting probe region overlaps with a LINE element. *Epn* expression was detectable by FANTOM5 CAGE expression. B, *Epn* expression upon 2i withdrawal in cells fractionated based on Rex1-GFP expression. 25hH and 25hL: Rex1-GFP high and low cells respectively sorted at 25 hours post 2i withdrawal. C, Sequence alignment of Exon3 of *Epn* and a rat EST transcribed from the syntenic region. *Epn* EST, mouse AK131952; rat EST, CA504619. D, Northern blot of wild type and *Epn* KO cells in 2i/LIF and 24 hours post 2i/LIF withdrawal. * non-specific hybridisation. 28s and 18S gel electrophoresis served as loading control. E, 5’RACE sequence confirming the 5’ start of *Epn* RNA. F, 3’RACE sequence confirming the 3’ end of *Epn* RNA. G, *Epn* expression relative to *β-actin* measured by RT-qPCR upon 2i/LIF component addition to LIF/serum (LS) culture. PD or/and CH were added to cells maintained in LIF/serum for indicated time. Mean+/-SD, n=3.

**Fig1 - Figure Supplement 2:**
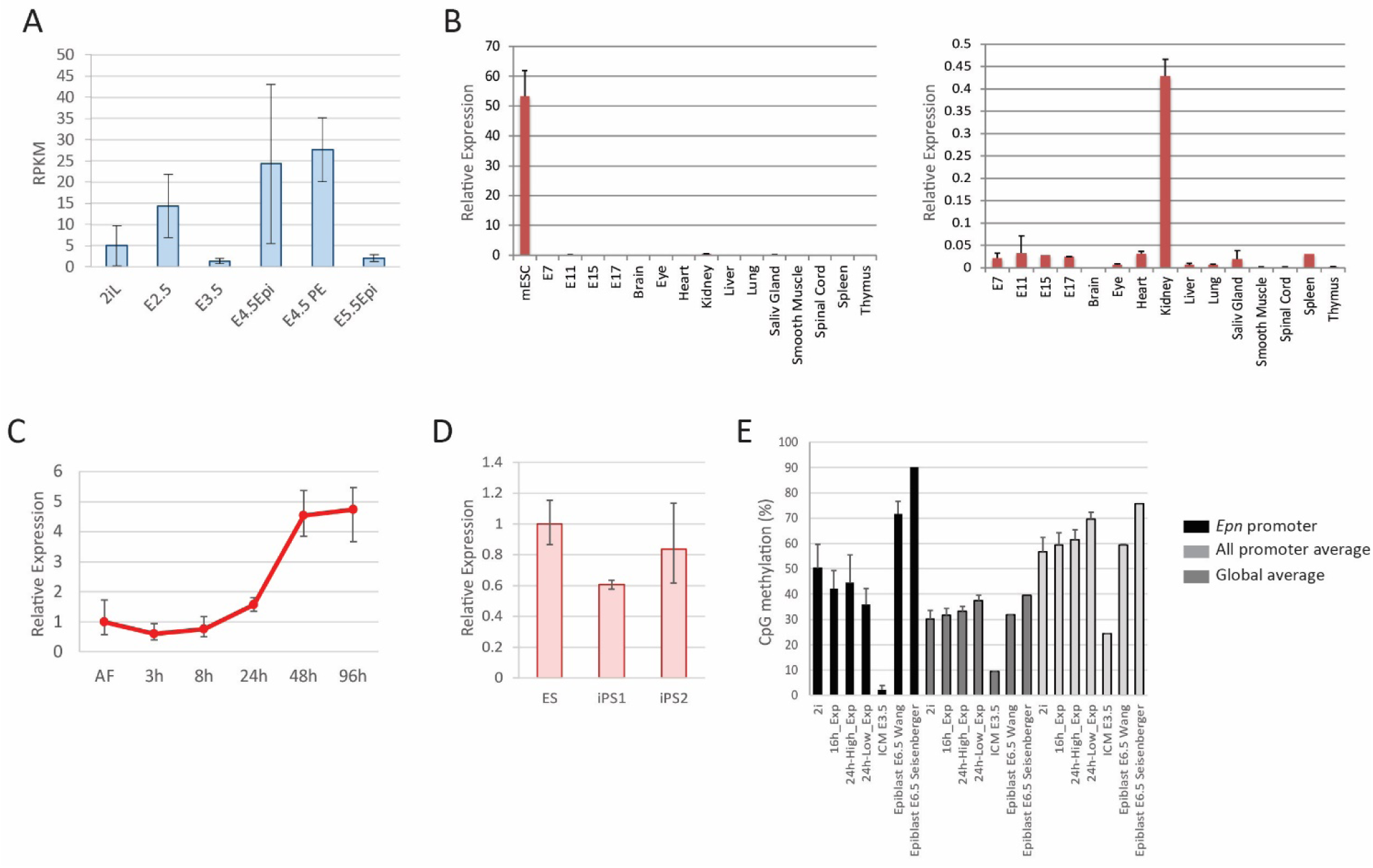
*Epn* expression and promoter methylation. A, *Epn* expression in early embryo development based on data from Boroviak et al 2015. Mean+/- S.E. n=3. B, *Epn* expression in later stages of embryo development and a panel of somatic tissues relative to *Hprt*. Mean+/-SD, n=3. C, *Epn* expression during EpiSC resetting. Mean+/-SD, n=3. D, *Epn* expression in established iPS clones. Mean+/-SD, n=3. E, Percentage of CpG methylation at *Epn* promoter, average of all promoters, and genome average upon 2i withdrawal and during early embryo development. Data are from Kalkan et al 2017, Seisenberg et al 2012, and Wang et al 2014.

**Fig2 - Figure Supplement 1:**
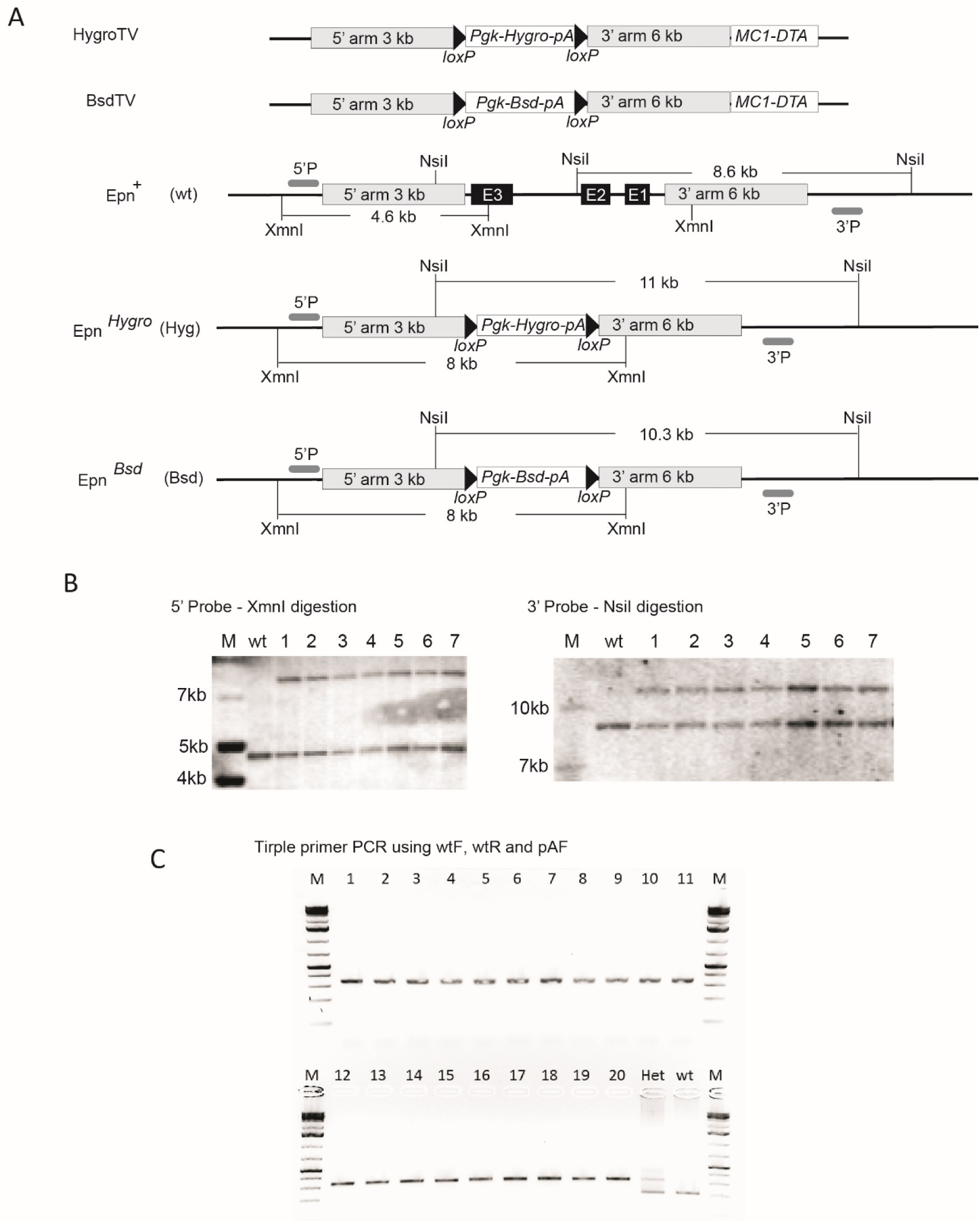
Generation of *Epn* KO ESCs. A, Targeting vector, gene targeting and Southern blotting validation strategies. B, Southern blotting confirming knockout of the first allele using an external probe. C, Triple primer PCR confirming ESC clones with both alleles targeted. Wild type (wt) and single allele targeted (Het) ESCs were used as PCR controls. M, molecular marker. Primer sequences are shown in the Supplementary file 1C.

**Fig2 – Figure Supplement 2:**
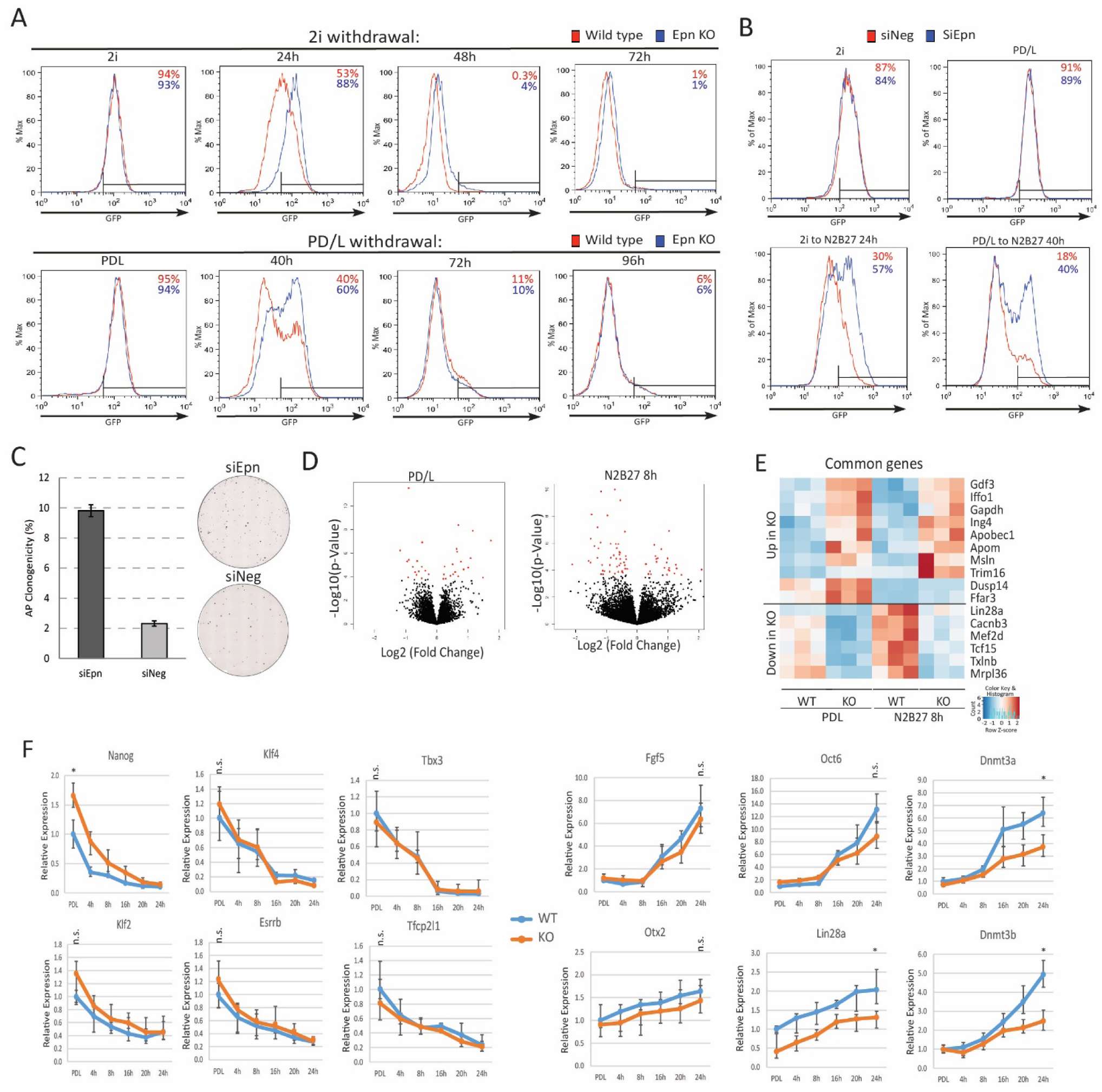
Phenotypic and molecular characterisation of *Epn* KO during naïve state exit. A, Rex1-GFP flow cytometry profile over 2i and PD/LIF withdrawal time course for wild type and *Epn* KO cells. B, Rex1-GFP flow cytometry profile of *Epn* knockdown in 2i and PD/LIF and after 24 hours and 40 hours in N2B27 respectively. C, Colony formation capacity of *Epn* knockdown cells 40 hours post PD/LIF withdrawal. Mean+/- SD, n=3. Representative AP staining images are shown. D, Volcano plots of differentially expressed genes in *Epn* KO compared to wild type ESC in PD/LIF and 8 hours after PD/LIF withdrawal. Red dots, statistically significant genes (Benjamini-Hochberg adjusted p<0.05) based on 3 independent wild type and KO lines. E, Significantly differential expressed genes common in both PD/LIF and 8h N2B27 conditions with fold change >1.5 or <0.7. F, Expression relative to *β-actin* of core naïve pluripotency factor network and peri-implantation epiblast genes in wild type and *Epn* KO ESCs upon PD/LIF withdrawal. Mean+/-SD, n=3. * p<0.05, student t-test; n.s. not statistically significant.

**Fig2- Figure Supplement 3:**
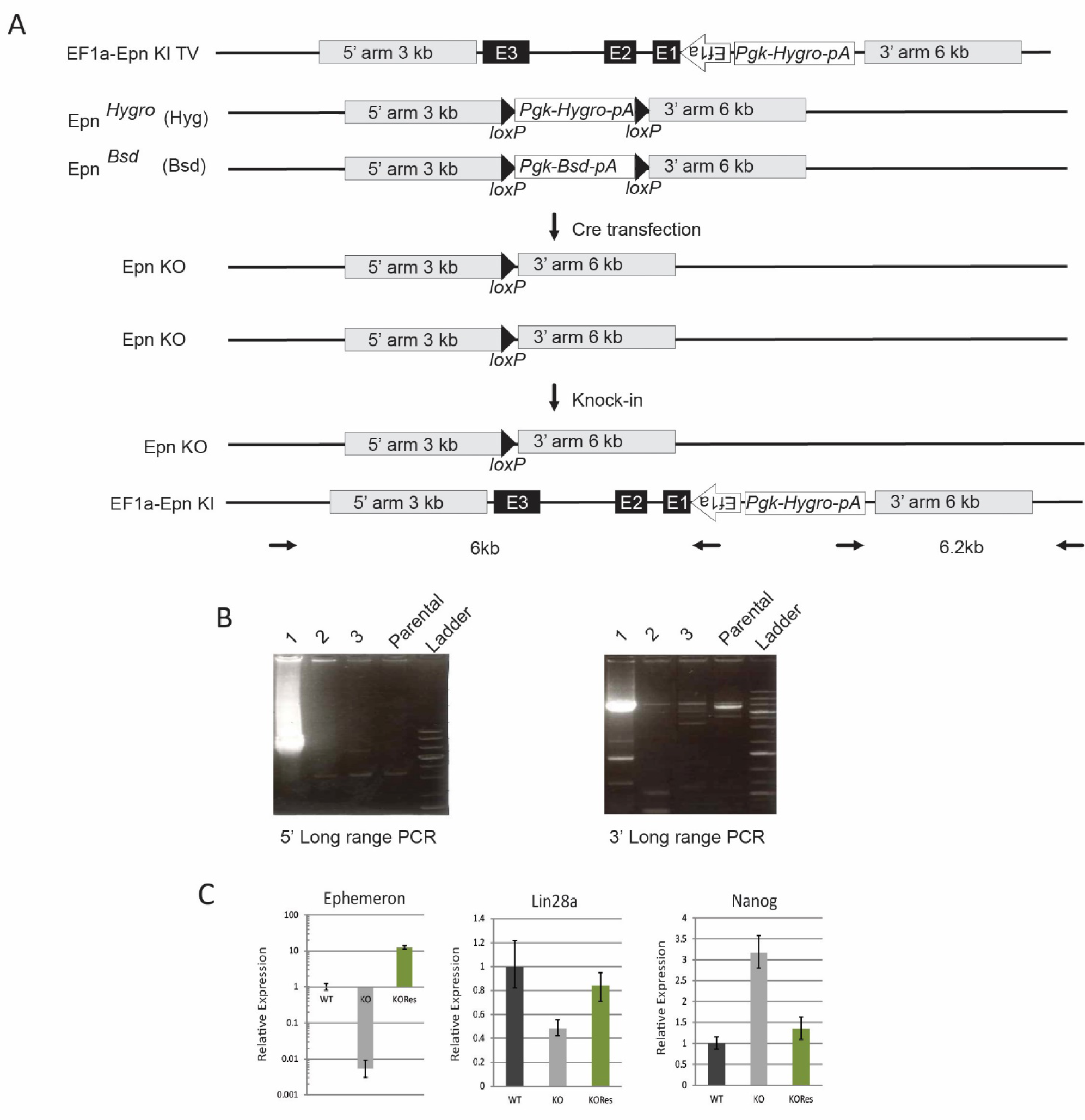
Generation of *Ephemeron* KO rescue ESCs. A, generation of *Epn* KO rescue cells by knock-in of the human *EF1a* promoter driving the genomic region containing all *Epn* exons. The *EF1a* promoter TSS site was cloned to be directly upstream of the first base of *Epn* exon 1. B, Long-range PCR genotyping of targeted clone. 5’ long range PCR product, 6 kb; 3’ long range PCR product, 6.2 kb. C, Expression of *Epn*, *Lin28a* and *Nanog* relative to *β-actin* in PD/LIF measured by RT-qPCR. Mean+/-SD, n=3.

**Fig2- Figure Supplement 4:**
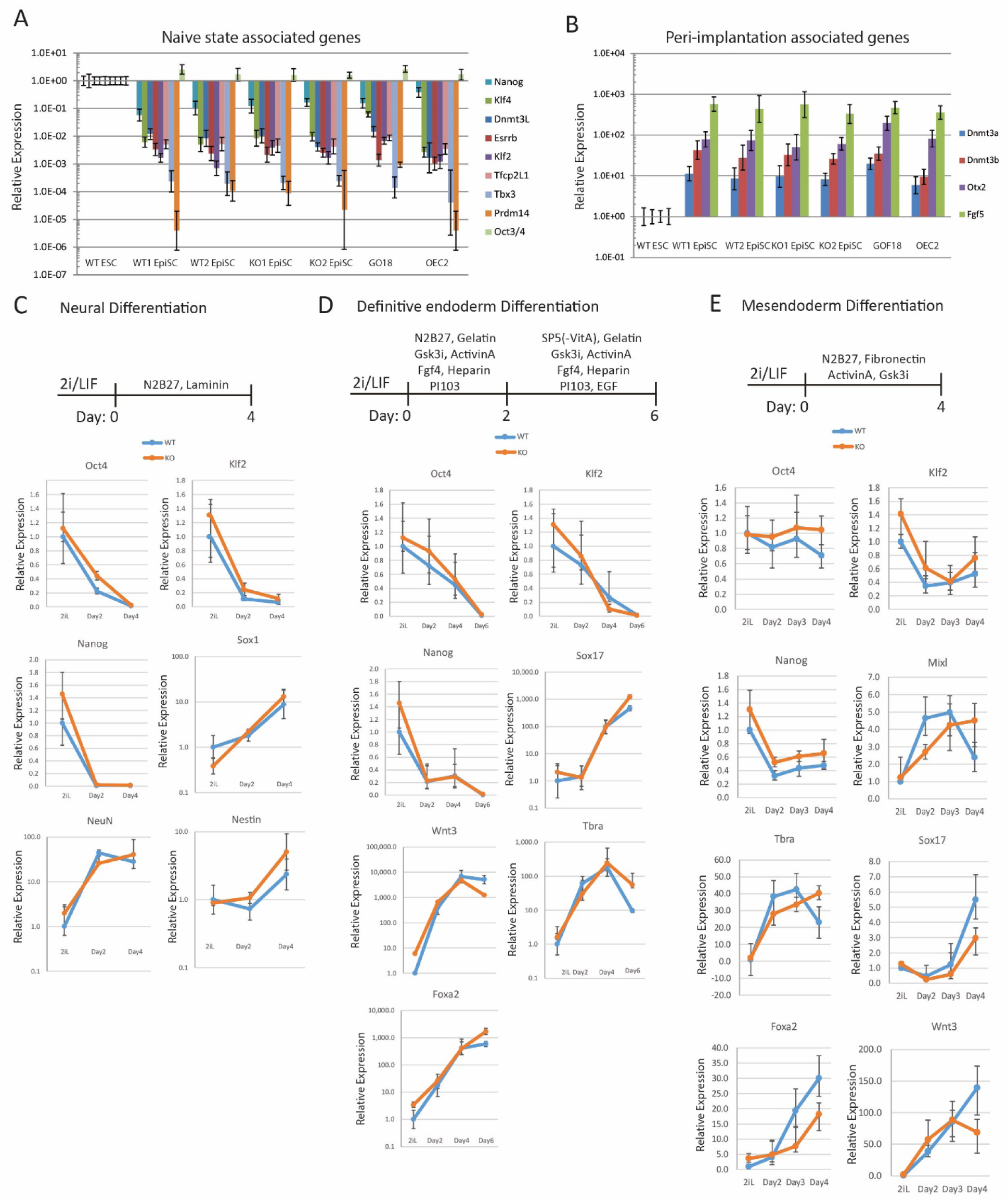
Differentiation capacity of *Epn* KO ESCs. A,B, naïve associated (A) and early peri-implantation epiblast associated (B) gene expression relative to *β-actin* in wild type and *Epn* KO ESC derived EpiSCs. Mean+/-SD, n=3. C, Neuronal differentiation protocol and gene expression of wild type and *Epn* KO ESCs relative to *β-actin*. Mean+/-SD, n=3. D, Definitive endoderm differentiation protocol and gene expression of wild type and *Epn* KO ESCs relative to *β-actin*. Mean+/-SD, n=3. E, Mesendoderm differentiation protocol and gene expression of wild type and *Epn* KO ESCs relative to β-actin. Mean+/-SD, n=3.

**Fig3 – Figure Supplement 1:**
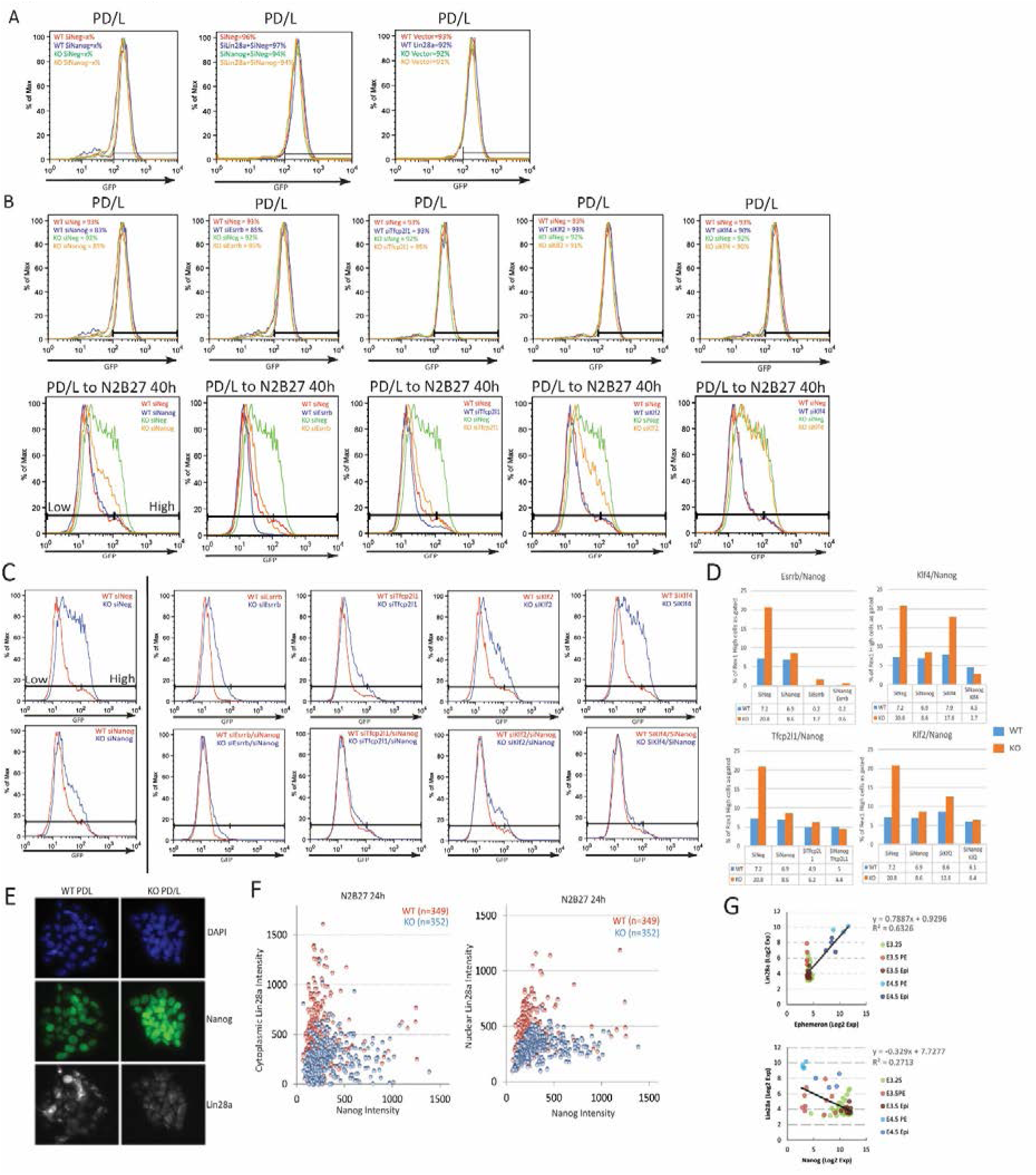
Characterisation of *Epn*, Nanog and Lin28a genetic interaction. A, Rex1-GFP profiles of indicated genetic manipulations of wild type and *Epn* KO cells in PD/LIF and 40 hours post PD/LIF withdrawal, related to Fig3A-C. B, Knockdown effects of several core pluripotency network factors on Rex1-GFP profile in wild type and *Epn* KO cells in PD/LIF and upon 40 hours withdrawal. C, Rex1-GFP profiles upon knockdown of core pluripotency factors singly or together with Nanog in wild type and *Epn* KO cells at 40 hours post PD/LIF withdrawal. D, Quantification of Rex1 high population as gated in C in wild type and *Epn* KO cells. E, Representative immunostaining images of wild type and *Epn* KO cells cultured in PD/LIF. F, Correlation analysis of Nanog and Lin28a protein expression based on quantification of fluorescent intensity of individual cells in wild type and *Epn* KO cells 24 hours post PD/LIF withdrawal. G, Correlation analysis of *Epn*, *Lin28a* and *Nanog* expression at single cell levels during pre-implantation embryo development based on published dataset (Ohnishi et al., 2014).

**Fig3 – Figure Supplement 2:**
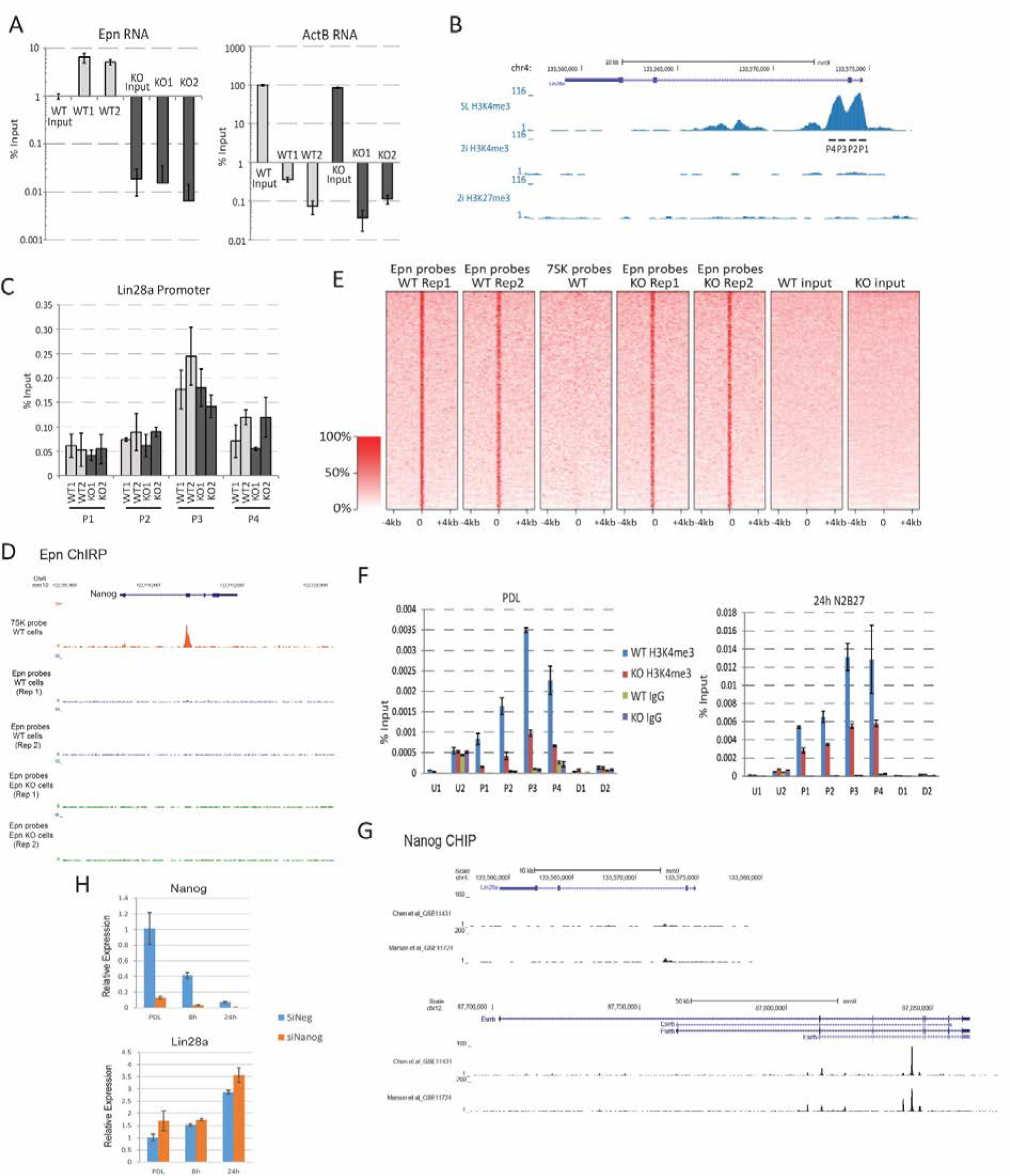
*Epn* does not act on chromatin. A, ChIRP using probe sets recognising *Epn* specifically enriched *Epn* RNA quantified by RT-qPCR. Mean+/-SD, n=3. B, *Lin28a* promoter region and H3K4me3 modification status base on published dataset (Marks et al., 2012). SL, serum/LIF. C, ChIRP using *Epn* probe set did not enrich *Lin28a* promoter chromatin quantified by qPCR. Mean+/-SD, n=3. The regions analysed are show in B. D, *Epn* ChIRP profile at *Nanog* locus. E, Genome-wide *Epn* ChIRP peaks in wild type and *Epn* KO cells. 7SK ChIRP probe was used as control. F, Lin28a promoter region H3K4me3 modification in wild type and *Epn* KO cells in PD/LIF and 24 hours post PD/LIF withdrawal. Mean+/-SD, n=3. G, Nanog ChIP profile at *Lin28a* and *Esrrb* loci based on published datasets (Marson et al 2008 and Chen et al 2008). H, *Lin28a* expression upon Nanog knockdown in PD/LIF and up to 24 hours post PD/LIF withdrawal. Mean+/-S.D, n=3.

**Fig4 – Figure Supplement 1:**
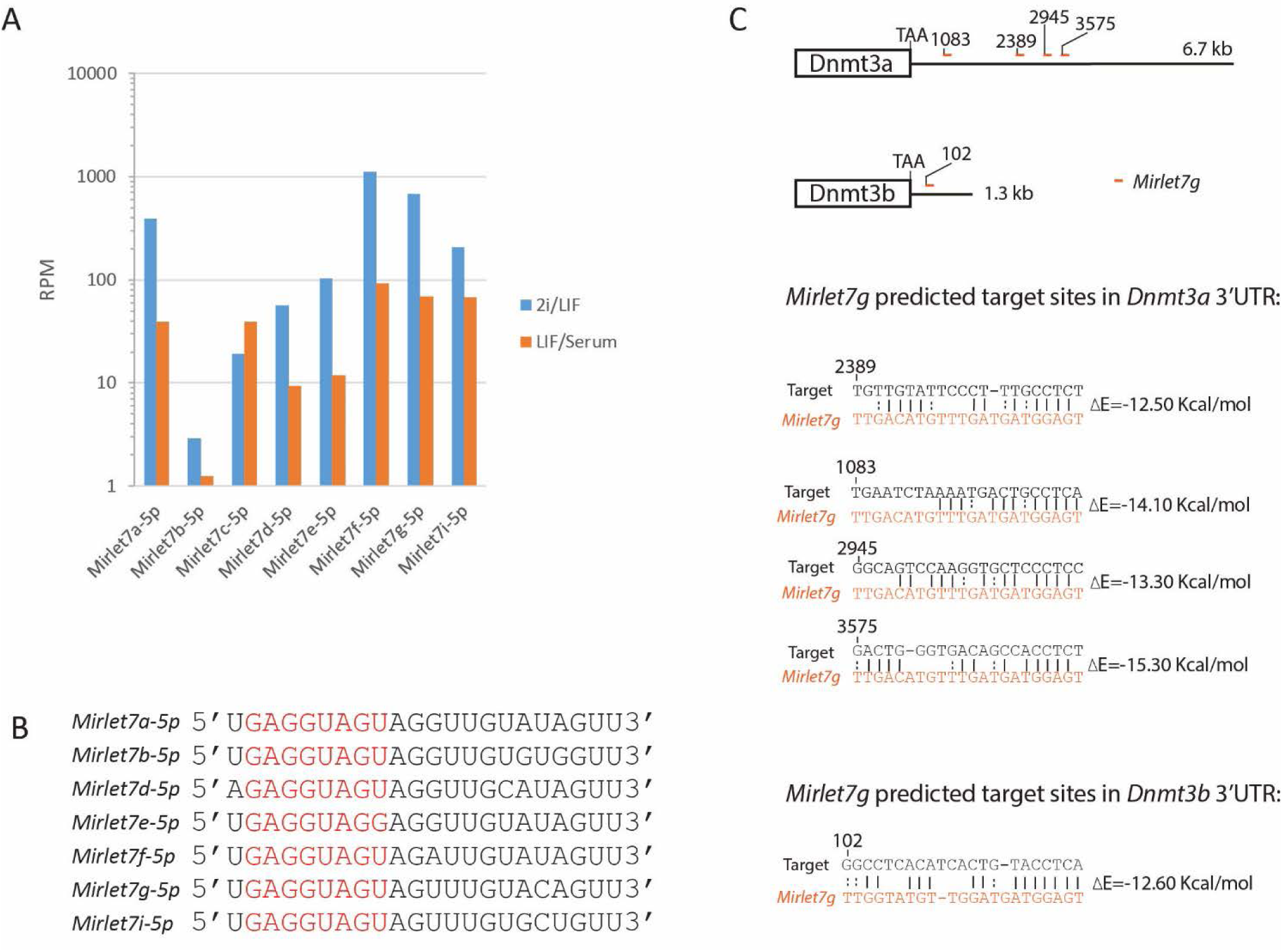
*Mirlet7* family mature miRNA expression, sequence and predicted *Mirlet7g* sites in the 3’UTRs of *Dnmt3a* and *Dnmt3b*. A, *Mirlet7* family mature miRNA expression in ESCs cultured in 2i/LIF and Serum/LIF from published dataset (Pandolfini et al., 2016). B, Mature miRNA sequence of Lin28a regulated *Mirlet7* members. Seed sequences are coloured in red. C, Predicted *Mirlet7g* target sites and target/miRNA heteroduplex structures and folding energy based on RNA22 V2 predictions.

**Fig5 – Figure Supplement 1:**
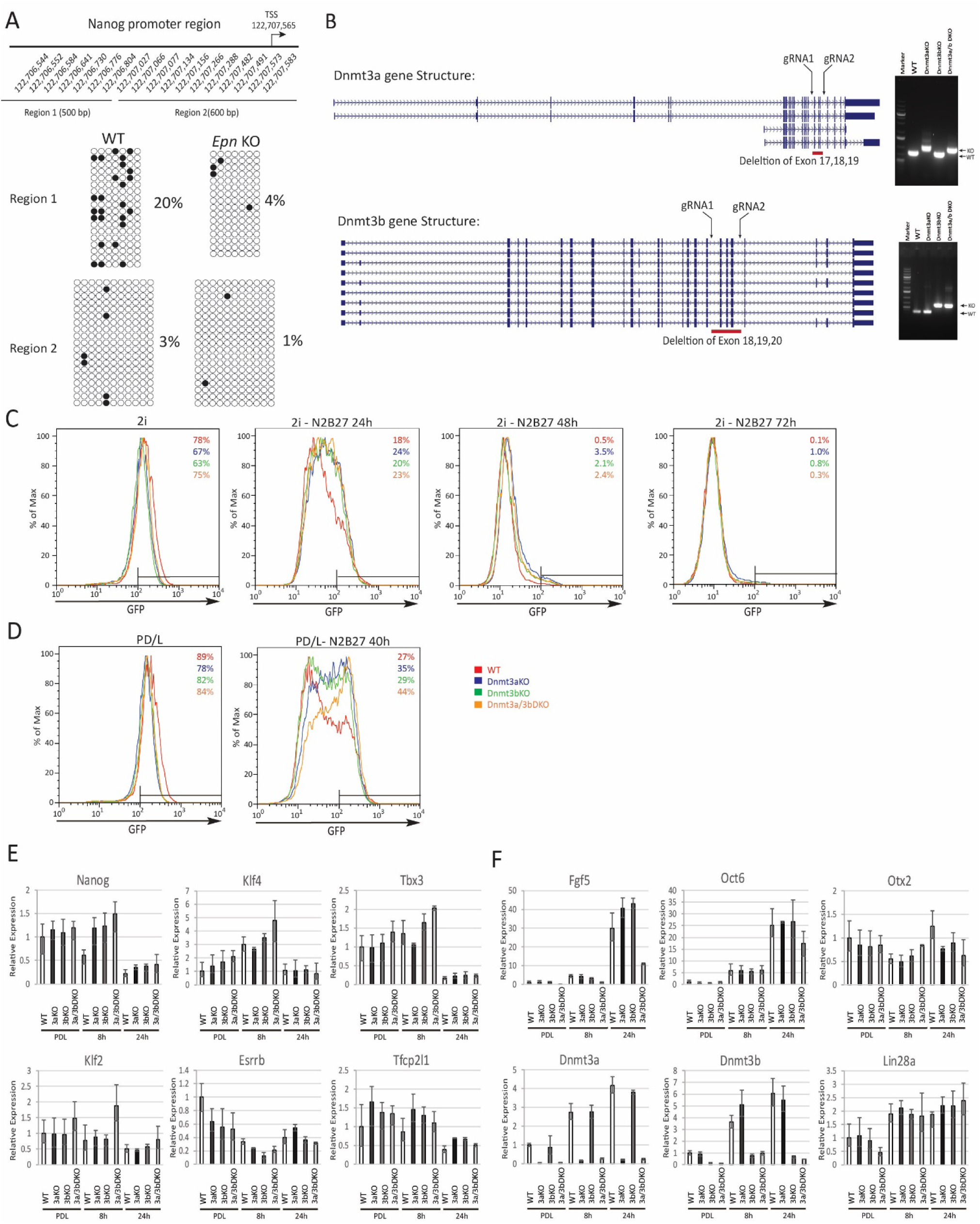
Phenotypic and molecular characterisation of *Dnmt3a/3b* KO in naïve state exit. A, Bisulfite sequencing analysis of Nanog proximal promoter CpG DNA methylation in wild type and *Epn* KO cells at 40 hours post PD/LIF withdrawal. The positions of cytosines (mm10) analysed are indicated. Black and white circles represent methylated and unmethylated cytosine. B, *Dnmt3a* and *Dnmt3b* variants with designed gRNA positions. Right panel, genotyping confirmation of homozygous deletions for *Dnmt3a* and *Dnmt3b* single and compound KO ESCs and the primer sequences are shown in Supplementary file 2B. C, Rex1-GFP profiles over 2i withdrawal time course for wild type and *Dnmt3a/3b* single and compound KO ESCs. Percentage of Rex1-GFP high cells are quantified. D, Rex1-GFP profiles for *Dnmt3a/3b* single and compound KO in PD/LIF and 40 hours post PD/LIF withdrawal. Percentage of Rex1-GFP high cells are quantified. E, Expression of core pluripotency factors relative to *β-actin* in *Dnmt3a/3b* single and dual KO ESCs upon PD/LIF withdraw. Mean+/- SD, n=3. F, Expression of peri-implantation markers relative to *β-actin* in *Dnmt3/3b* single and dual KO ESCs upon PD/LIF withdrawal. Mean+/-SD, n=3.

